# Epistemic uncertainty challenges aging clock reliability in predicting rejuvenation effects

**DOI:** 10.1101/2023.12.01.569529

**Authors:** Dmitrii Kriukov, Ekaterina Kuzmina, Evgeniy Efimov, Dmitry V. Dylov, Ekaterina E. Khrameeva

**Affiliations:** Skolkovo Institute of Science and Technology, Moscow, Russia; Artificial Intelligence Research Institute, Moscow, Russia

**Keywords:** Rejuvenation, cell reprogramming, epistemic uncertainty, epigenetic aging clocks, dataset shift, DNA methylation

## Abstract

Epigenetic aging clocks have been widely used to validate rejuvenation effects during cellular reprogramming. However, these predictions are unverifiable because the true biological age of reprogrammed cells remains unknown. We present an analytical framework to consider rejuvenation predictions from the uncertainty perspective. Our analysis reveals that the DNA methylation profiles across reprogramming are poorly represented in the aging data used to train clock models, thus introducing high epistemic uncertainty in age estimations. Moreover, predictions of different published clocks are inconsistent, with some even suggesting zero or negative rejuvenation. While not questioning the possibility of age reversal, we show that the high clock uncertainty challenges the reliability of rejuvenation effects observed during in vitro reprogramming before pluripotency and throughout embryogenesis. Conversely, our method reveals a significant age increase after in vivo reprogramming. We recommend including uncertainty estimation in future aging clock models to avoid the risk of misinterpreting the results of biological age prediction.

## Introduction

Reprogramming aged somatic cells into pluripotency or other progenitor states was consistently shown to ameliorate various aging-associated features, either by applying different transcription factors (TFs) or by introducing small molecules to cells in vitro or in vivo [1–5]. To quantify the effect of age reversal, researchers resort to various methods, including “epigenetic aging clock” models built from DNA methylation (DNAm) data with diverse machine learning (ML) approaches. These approaches are widely employed to compare the “biological age” of reprogrammed cells to that of control cells [6] (Figure **1**a,b). These clocks are easy to use and can assess aging from the organismal to the cellular levels, which is particularly valuable in experiments where the large-scale parameters of organismal aging cannot be measured, such as *in vitro* cellular reprogramming. Notably, a similar approach has recently been used to demonstrate rejuvenation in the course of embryonic development [7–9].

**Figure 1:**
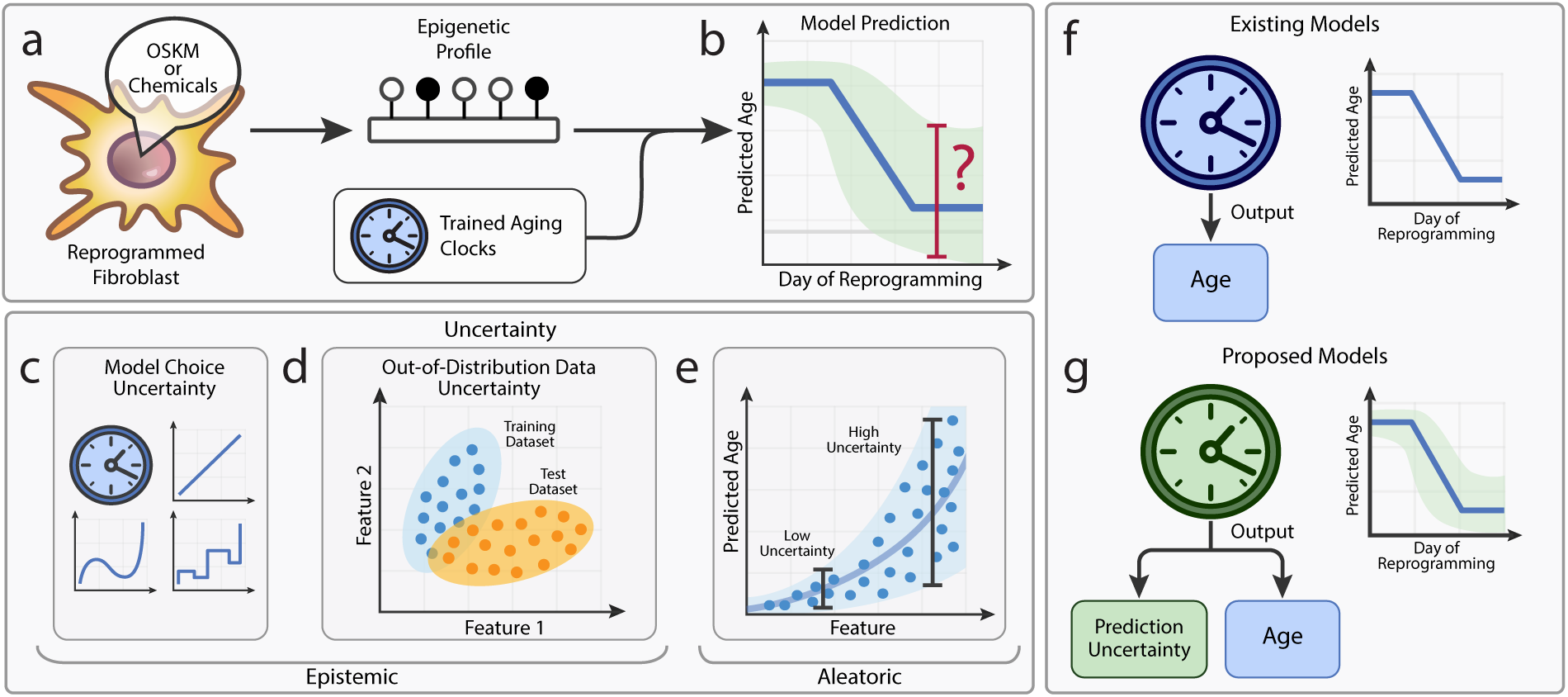
Prediction uncertainty is an essential component for clinically relevant aging clocks. **a**, A common pipeline for testing rejuvenation effect using epigenetic aging clocks. **b**, Current aging clocks estimate biological age during the reprogramming process; however, they lack epistemic uncertainty quantification. **c**, By selecting a model to make predictions from data, a researcher implicitly introduces model uncertainty. **d**, Out-of-distribution uncertainty arises when the testing data samples are not represented in the training distribution. **e**, Aleatoric uncertainty comes from the intrinsic variability in data, *e.g.* when the same level of a feature corresponds to different ages. **f**, None of the existing aging clocks estimates epistemic uncertainty. **g**, We propose to use aging clocks capable of predicting uncertainty, which could mitigate the potentially erroneous effects of clock predictions on clinical decision-making.

The primary assumption of aging clocks is that the deviation Δ of predicted age from the chronological age *C* represents an accelerated or decelerated aging, that is, an increase or decrease in the biological age *B* [10, 11]. One can express it as *B* = *C* +Δ. Since biological age cannot be measured directly (*i.e.*, it lacks a definitive “ground truth”), the epigenetic age estimated by the clocks is therefore considered a proxy measure of the biological age [12]. Consequently, owing to the fact that DNAm patterns are strong indicators of past influences and future health outcomes, the epigenetic age is proposed to serve as a biomarker for measuring the effects of pro-longevity interventions in clinical trials [10, 13, 14].

However, before aging clocks could be integrated into clinical practice, these models should provide an estimate of uncertainty for their own predictions. Uncertainty manifests itself in three ways [15, 16]: (i) Model choice uncertainty, part of a broader category known as epistemic uncertainty, represents how well a proposed model (its architecture, parameters, metrics, *etc.*) reflects the actual underlying process (Figure **1**c). (ii) Out-of-distribution (OOD) uncertainty, another type of epistemic uncertainty, emerges when the testing data are not represented in the training data distribution, leading to a high risk of model prediction failure (Figure **1**d). (iii) Aleatoric uncertainty originates from data variations that cannot be reduced to zero by the model (*e.g.*, when the same DNAm level corresponds to different ages) (Figure **1**e).

Dataset shift [17] term describes the case of OOD sampling where the testing population is under-represented within the training distribution. Dataset shift can be decomposed into covariate shift (*e.g.*, differently distributed DNAm values) and response shift (*e.g.*, different age ranges). Notably, batch effect serves as a notorious example of dataset shift in the field of omics data analysis because it indicates differences in covariate or response distributions between sample groups.

From the clinical perspective, epistemic uncertainty must be estimated to make reliable conclusions about whether to trust a model or not [15]. Specifically, epistemic uncertainty resulting from the dataset shift should be scrutinized, considering the prevalence of batch effects in biological data [18, 19]. However, most popular DNAm aging clocks fail to meet this criterion (Figure **1**f) because they are typically built using algorithms from the penalized multivariate linear regression (MLR) family [20] (*e.g.*, ElasticNet). Such algorithms do not yield information on any of the uncertainties, except for the error between chronological and predicted ages in the training data (*e.g.*, mean or median absolute errors, *MAE* or *MedAE*).

In this work, we question the applicability of existing aging clock methodology for measuring rejuvenation by specifically examining prediction uncertainty (Figure **1**). To achieve this, we reanalyze published data of putative rejuvenation, focusing on extreme cases of anticipated dataset shift, such as cellular reprogramming and embryonic development.

Because biological age measurements cannot be verified explicitly, we introduce four different indirect approaches to this problem: (i) Elucidating covariate shift in DNAm values between the datasets of aging and rejuvenation. (ii) Evaluating the agreement between different aging clocks in predicting rejuvenation. (iii) Exploring if rejuvenation datasets can be employed reciprocally to predict normal aging. (iv) Assessing whether an aging clock capable of estimating its own uncertainty (Figure **1**g) would demonstrate a significant age reversal in putative rejuvenation experiments.

We propose a comprehensive framework for implementing these approaches. By leveraging this framework, we aim to elucidate the most critical limitations in applying aging clock models to rejuvenation studies, which should be solved in order to drive broader acceptance of these models within the longevity community.

## Results

### Covariate shift can lead to biologically meaningless predictions

To introduce the concept of covariate shift in the field of aging clocks, we first explored a simple, low-dimensional example. In particular, we used two parameters (biomarkers) to construct an elementary aging clock for predicting chronological age in humans: weight and height (Figure **2**a; see Methods). These two biomarkers strongly correlate with age during the first twenty years of human life [21], therefore they can legitimately be used for age prediction. We analyzed the data of body measurements for male humans ranging from 1 to 25 years old (an approximate end of body growth), including both healthy controls [21] and individuals with the achondroplasia disorder [22, 23] typically characterized by a shorter length of arms and legs [24].

**Figure 2:**
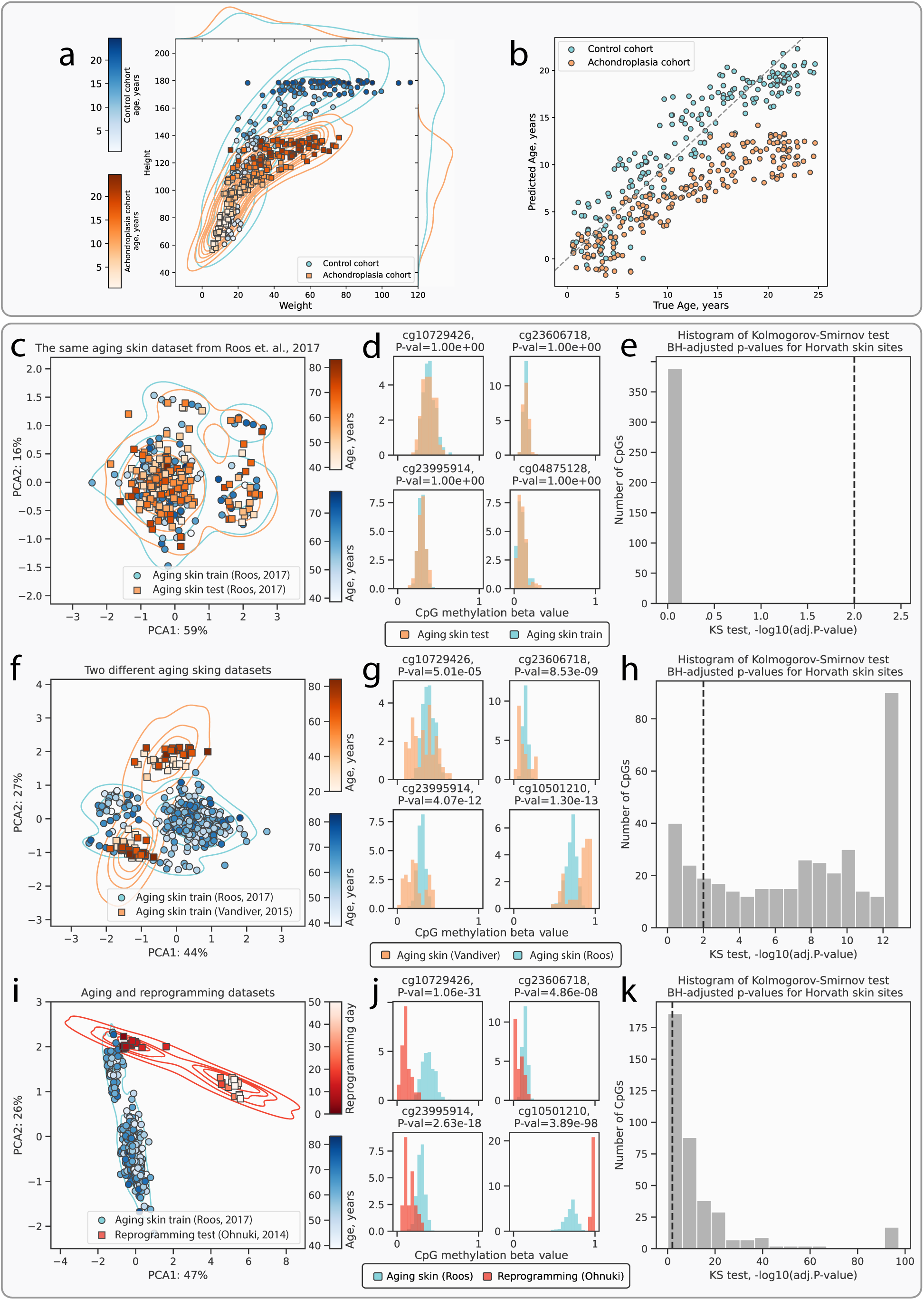
Identification of covariate shift and its impact on aging clock models. **a**, An example of weight and height covariate shift between the control and the achondroplasia cohorts. **b**, Ages predicted by a model trained on the weight and height measurements of the control cohort. The predictions are significantly biased for the achondroplasia cohort, which is a purely technical phenomenon caused by shifted covariates. **c,f,i**, Principal component analysis (PCA) of DNAm samples shows no covariate shift between the training and testing splits of the same aging skin dataset [28] **(c)**, a moderate covariate shift between the different aging skin datasets [28, 29] **(f)**, and a strong covariate shift between the aging skin dataset [28] and the in vitro fibroblast reprogramming dataset [30] **(i)**. **d,g,j**, Histograms of beta values for top-4 age-correlated CpG sites demonstrate no shifts between the two subsets of data **(d)**, moderate shifts between the aging skin datasets from different studies **(g)**, and strong shifts between the aging skin and the reprogramming datasets **(j)**. **e,h,i**, Histograms of *−log*_10_(adj. *P* values) illustrate that no DNAm sites were rejected by the KS two-sample test at the 0.01 significance level (see Methods) confirming the absence of covariate shift **(e)**, 81% of DNAm sites were rejected by the KS test confirming the presence of moderate covariate shift **(h)**, and 86% of DNAm sites were rejected by the KS test confirming the presence of strong covariate shift **(k)**. Percentages on the axes in **c**, **f**, **i** indicate the amount of variance explained by the corresponding principal components. Representative sites for histograms **d**, **g**, **j** were chosen from the top-four sites ordered by their correlation with chronological age.

Inspired by the common framework of aging clocks construction established in earlier works [25–27], we trained a multivariate linear regression (MLR) model using body measurements of a cohort of healthy individuals to predict their chronological age, which yielded a good performance on the training data (*MAE*=2.3 years, *R*^2^=0.84). For the achondroplasia cohort, these clocks predicted consistently lower ages (Figure **2**b), which would be viewed as decelerated aging in the context of other aging clocks. However, this interpretation has no biological support because the average lifespan of people with achondroplasia is around 10 years shorter than that of control individuals due to early-life mortality [24]. We assume that this underestimation of ages in the achondroplasia cohort is caused by a covariate shift in the analyzed data leading to a huge OOD uncertainty, which is not taken into account by the model. Indeed, the distributions of covariates (weight and height) differs significantly between the training and the testing data (Kolmogorov-Smirnov (KS) test, *P* value<0.0003; Figure **2**a). In general, any significant differences between the observed distributions in training and testing samples should caution us against the casual application of ML models.

Next, we expanded our analysis beyond this simplistic case to examine various DNA methylation datasets and to thoroughly investigate possible covariate shifts in the context of epigenetic aging clocks.

### Reprogramming and embryogenesis datasets exhibit significant covariate shifts relative to aging datasets

Covariate shifts can arise from various intrinsic and technical factors, including differences in sampling sources and locations, tissue cell content, sample handling techniques, instrumental effects, *etc.* [18]. To estimate the extent of covariate shift in DNA methylation (DNAm) studies, we analyzed four representative scenarios in the order of increasing expected difference between the distributions of DNAm patterns: (i) One aging dataset split into two subsets; (ii) Two independent aging datasets; (iii) Aging *vs.*cellular reprogramming in vitro and in vivo; and (iv) Aging *vs.*early embryogenesis, for which epigenetic rejuvenation was also demonstrated [7].

As our study was aimed at testing the analytical framework rather than at providing comprehensive coverage of dataset shift in all existing data of putative rejuvenation, we focused on several outstanding cases of reprogramming, where DNAm was profiled across as many time points as possible, and the datasets were published in open access (see Methods).

First, we examined a DNAm dataset of aging human skin by Roos et al. [28] split randomly into the training and testing subsets. As anticipated, we detected no discernible covariate shift: the subsets are indistinguishable by the PCA (Figure **2**c), DNAm value distributions of at least top-4 age-correlated CpG sites (CpGs) from both subsets perfectly overlay each other (Figure **2**d), and the KS test for distribution similarity further confirms the lack of covariate shift (Figure **2**e).

Second, for two independent datasets of aging human skin [28, 29], moderate covariate shift is evident from a similar analysis (Figure **2**f,g), with the KS test indicating substantial differences in individual distributions (Figure **2**h): 81% percent of sites were rejected by the test (meaning that these CpGs have different distributions). On the other hand, a joint analysis of two aging mouse liver datasets [31, 32] displays minimal covariate shift (only 1% of CpGs are rejected) (Figure **S1**d-f).

Third, a comparison of the aging human skin dataset [28] with two datasets of in vitro human fibroblast reprogramming [4, 30] revealed strong covariate shifts: at the early stages, the fibroblasts appear to closely resemble aging skin samples in their principal component (PC) coordinates (Figure **2**i; Figure **S1**a), but, as the reprogramming progresses through the maturation phase, a notable departure from the aging skin samples can be observed (86% and 69% of CpGs are rejected, respectively; Figure **2**j,k; Figure **S1**b,c). Such sample behavior during in vitro reprogramming might suggest that the reprogrammed cells acquire some phenotype unobservable in vivo.

To shed more light on this hypothesis, we further compared a dataset of in vivo reprogramming in mouse liver [3] with merged liver samples from two datasets of mouse aging [31, 32], as these two studies demonstrated no significant difference in the previous analysis. As a result, we detected a moderate covariate shift according to the PCA and the KS test (32% of CpGs were rejected, Figure **S1**g-i), which might imply that the in vivo conditions are better at preserving the normal phenotypic characteristics.

Fourth, to address the “ground zero” hypothesis of epigenetic rejuvenation during embryogenesis [33], we compared mouse embryos [34] with blood aging samples [31], as both of these datasets were used previously to demonstrate this phenomenon [7]. The first stages of embryogenesis strongly diverge from the aging samples, while the later stages align closer to the aging cluster on PCA, suggesting a moderate covariate shift, which is further supported by the KS test (15% of CpGs were rejected, Figure **S1**j-l).

These results collectively suggest that the DNAm covariates can lead to significant shifts between datasets, thereby implicitly increasing the risk of clock prediction failure. Given these risks, we advocate for the routine checks of covariate shifts between datasets using the combination of PCA and KS test or other techniques reviewed elsewhere [17] before applying aging clocks.

### Aging clocks are inconsistent in their predictions for reprogramming-induced rejuvenation

When choosing a specific machine learning model to construct an aging clock, a researcher inevitably introduces model uncertainty (Figure **3**a). The choice of model family (*e.g.*, Linear Regression) includes certain assumptions about how the true process of epigenetic aging, not accessible *a priori*, works. Therefore, by training different aging clocks on different data or using different ML model families, one can expect to obtain varying clock CpG subsets [35] and poorly consistent predictions of normal aging [36].

**Figure 3:**
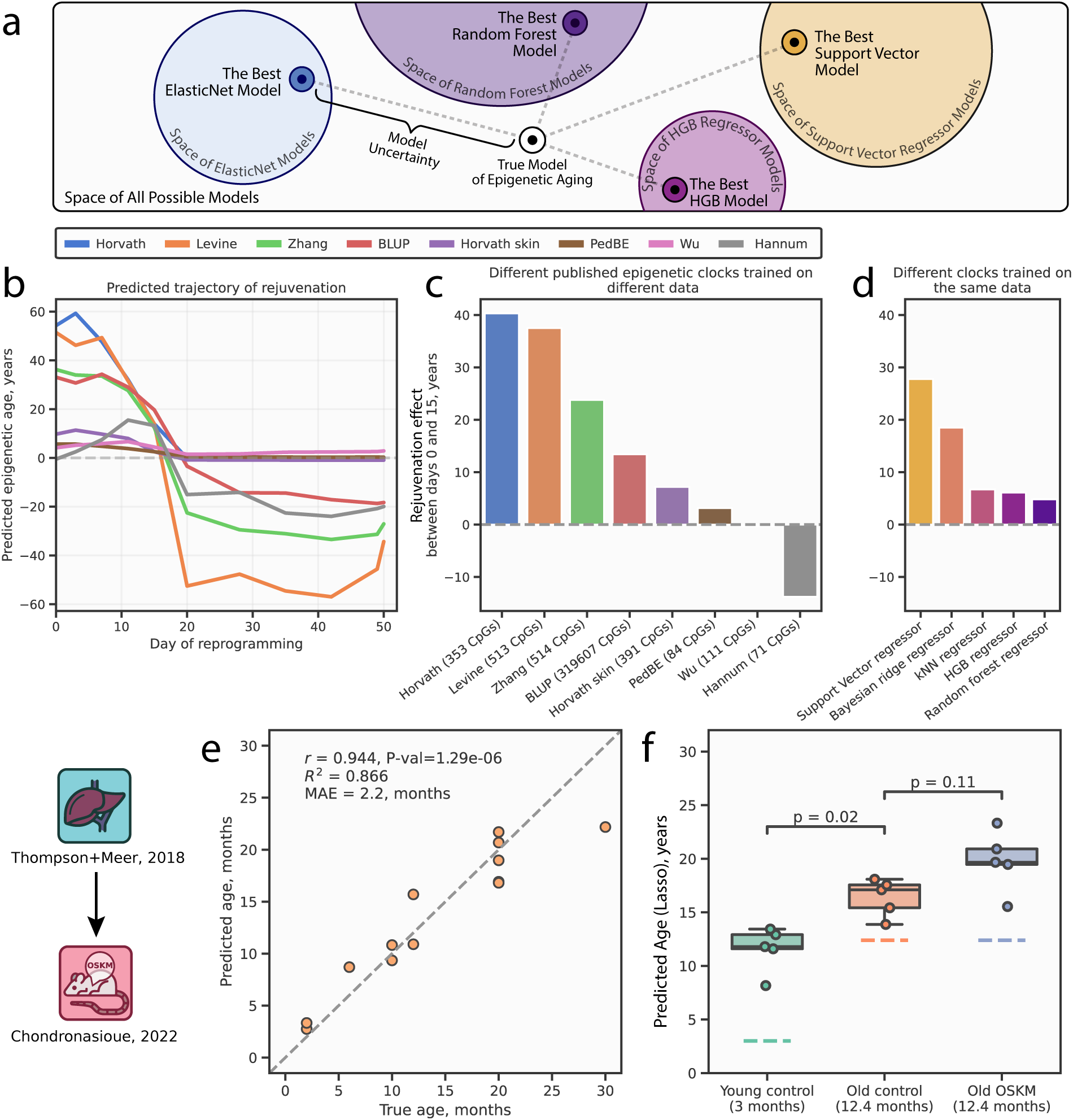
Inconsistency between the estimations of reprogramming-induced rejuvenation provided by the aging clocks. **a**, A schematic representation of model uncertainty stemming from the choice of a particular model. **b**, Published aging clocks (most of which are ElasticNet-based) trained on different DNAm datasets predict different trajectories of rejuvenation, as cellular reprogramming progresses. The dashed line represents zero epigenetic age. **c**, Aging clocks showing differences in the rejuvenation effect accumulated between reprogramming days 0 and 15, bar colors match line colors from **a** and represent different published clocks. Dashed line represents lack of changes in epigenetic age. **d**, Inconsistency of predictions holds for the clocks built using different ML model types and trained on the same dataset. Bar colors represent different models. The dashed line represents lack of changes in epigenetic age. **e**, Scatter plot of the de novo Lasso clock model performance on the testing subset (see Methods). Pearson’s correlation coefficient (*r*), the associated *P* value (*Pval*), *R*^2^ score (*R*^2^), and mean absolute error (*MAE*) are displayed. The dashed line corresponds to the points of equality between the predicted and chronological ages. Dots represent individual testing samples. **f**, Epigenetic ages predicted by the de novo Lasso clock for the in vivo liver reprogramming dataset [3] containing samples from the young (n=5, green boxplot and sample dots) and old (n=5, orange boxplot and sample dots) control mice, and old reprogrammed mice (n=5, blue boxplot and sample dots). Samples from the transiently reprogrammed mice exhibit insignificantly increased (*P* value=0.11, two-sided MWW test) epigenetic age in comparison to the control group. Dashed lines represent chronological ages for the respective cohorts. Blue icon indicates the training dataset, red icon indicates the reprogr7amming dataset. Detailed dataset descriptions are presented in Supplementary Tables 1 and 2.

To demonstrate a critical role of model uncertainty, we leveraged eight published aging clocks trained on different CpG sets and tissue types [26, 27, 37–41] and applied them to two in vitro reprogramming datasets (see Methods). As expected, all these clocks varied greatly in their predictions across the reprogramming timeline (Figure **3**b, Figure **S2**a). We further focused on the period of the first three weeks of reprogramming (from initiation to maturation), as the end of the third week (approximately, day 20) marks the loss of somatic identity and the increasing risk of teratoma formation [1], which is undesirable from the clinical standpoint. A comparison of differences between the ages estimated at the beginning and at the last available time point before the end of this period (that is, day 15 for Ohnuki et al. [30] and day 17 for Gill et al. [4]) exhibits evident inconsistencies for both datasets, ranging from the Horvath multi-tissue clock [26] predicting age reversal by 40 years to the Hannum blood clock [27] predicting age increase by 13 years (Figure **3**c and Figure **S2**a,b).

To test the hypothesis that these discrepancies might have arisen from the differences in training datasets rather than from the clock models themselves (most of which are based on ElasticNet regression), we trained different ML models on the same dataset of aging human skin [28] and discovered that their predictions of rejuvenation demonstrate a considerable instability as well (Figure **3**d, Figure **S2**c). Despite these inconsistencies, most models indicated epigenetic age reversal. Therefore, the clocks could potentially serve as qualitative (or binary) predictors of rejuvenation for the in vivo studies.

To explore this possibility, we estimated biological ages in the in vivo reprogramming dataset [3]. For that, we developed new aging clocks by fitting a Lasso regression over the combined dataset of aging mouse liver [31, 32] (see Methods) considering that the CpGs in these aging and reprogramming datasets poorly overlap with the existing clock models, thus impeding their use. Our Lasso clocks yielded quite robust performance (Figure **3**e), but they failed to register any rejuvenation in old mice treated with OSKM factors, compared to old control mice (Figure **3**f). Notably, all models consistently predicted higher ages for all control liver samples, suggesting a response shift [17] (*i.e.*, *P_train_*(*Y*) ≠ *P_test_*(*Y*)) between the training and testing datasets, which might result from the differences in DNAm patterns between the training and testing mouse strains (inbred C57BL/6 and transgenic i4F-B, respectively).

Taken together, our findings highlight a considerable instability of predictions with regard to the choice of both the training dataset and the model family employed for prediction. This lack of agreement between the aging clocks could, supposedly, result from the covariate and the response shifts described previously in this study, as well as from the manifestation of model uncertainty, leading to the divergent rejuvenation dynamics.

### Reprogramming data cannot be used to predict normal aging

We further examined the interchangeability of the aging and reprogramming datasets. When the regression models are trained, they need to satisfy a certain degree of correlation expressed, for example, as the *R*^2^ score or mean absolute error (*MAE*). Supposedly, in an ideal case, when a clock predicts age with absolute accuracy (*R*^2^=1.0, *MAE*=0), the predicted ages can be used interchangeably with the true chronological ages (because they are equal) to train another model on the testing data, and to predict ages in the original training dataset with the same accuracy. In reality, this ideal case is never observed due to the technical and biological variations in the training and testing samples and sampling techniques, and due to model under- or overfitting. But we hypothesize that if these effects are small (*i.e.*, if there is little epistemic uncertainty), then the reciprocal prediction of training data by the ages predicted for testing data should still be possible, albeit with some degree of error.

To evaluate this mutual interchangeability of datasets from the perspective of model training, we developed an Inverse Train-Test Procedure (ITTP, see Methods) (Figure **4**a,b). Considering the availability of ground truth measurements, we divided the ITTP use cases into two categories. In case 1, when comparing two aging datasets, the biological ages are available for the testing dataset *X_te_, Y_te_* (Figure **4**a) because they can be approximated by the chronological ages due to the fact that the chronological ages provide important information on the biological ones [42]. According to the ITTP, we first train model 1 (*e.g.*, a linear regression) on the training set *X_tr_, Y_tr_*to predict the ages of testing samples *Ŷ_te_*, where the hat symbol^denotes predicted values. Second, model 2 is trained using the testing features *X_te_* and the ages predicted by the model 1 *Ŷ_te_*. If the datasets are indeed interchangeable, similarly good performance metrics are expected for the predictions made by model 2 for the original training samples (Figure **4**a).

**Figure 4:**
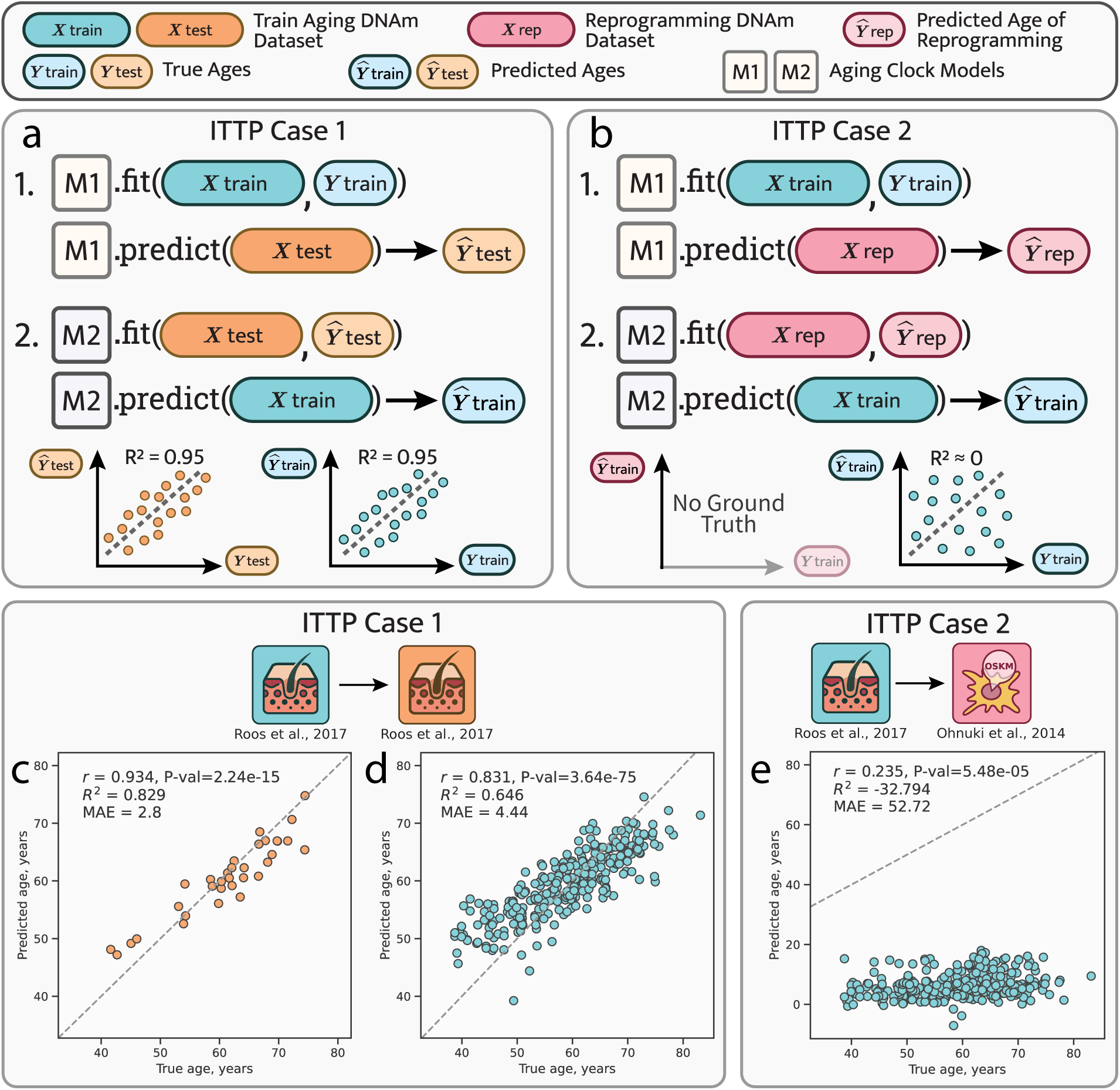
Inverse Train-Test Procedure (ITTP) demonstrates incorrect predictions of aging data based on reprogramming data. **a**, In ITTP case 1, where both datasets contain chronological age values (a verifiable case), interchangeability is evidenced by the comparable performance metrics across both procedural steps. **b**, ITTP case 2 is characterized by one dataset containing chronological age values and the other lacking them (an unverifiable case). Here, only the second-step performance metrics are computable. The failure of a hypothetical reprogramming dataset in the ITTP is indicated by the poor performance at the second step. **c,d**, Application of the ITTP to the aging human skin dataset [28] split into the training and the testing subsets as 90% and 10%, respectively (see Methods). As the performance metrics computed at both steps 1 (**c**) and 2 (**d**) are high, these datasets are considered interchangeable. **e**, Application of the ITTP to the aging human skin dataset [28] and the human fibroblast reprogramming dataset [30]. Poor performance metrics at step 2 qualify datasets as non-interchangeable — the reprogramming dataset cannot be used to predict aging data. Blue icons indicate the training datasets and orange icons indicate the testing dataset. The red icons indicate the reprogramming dataset. Detailed dataset descriptions are presented in Supplementary Tables 1 and 2.

In case 2, when leveraging an aging DNAm dataset {*X_tr_, Y_tr_*} and a reprogramming dataset {*X_rep_*}, we cannot approximate biological ages for the testing dataset (Figure **4**b) because we do not expect the biological age of reprogrammed cells to stay approximately the same as their chronological age, as would be the case for a dataset of normal aging. As before, we train model 1 on the training data and predict the ages of reprogramming samples *Ŷ_rep_*. In this scenario, we cannot validate our predictions due to the lack of ground truth values of biological age for reprogrammed cells, so we can only assume that the predictions are correct and use them to train model 2. If, as a result, we observe good performance metrics between the model 2 predictions and the ages of the original training dataset, then we can still assume interchangeability, regardless of the intermediary predictions of model 1. On the other hand, if we obtain poor prediction accuracy for model 2, then we infer that these datasets are not interchangeable, and that such prediction failure was supposedly caused by a large epistemic uncertainty, presumably originated from a substantial dataset shift. In summary, the ITTP approach contributes to our multifaceted estimation of prediction uncertainty in aging and reprogramming.

We further applied this procedure to all pairs of datasets described previously (Figure **2** and Figure **S1**) and used the Lasso regression model family as models 1 and 2 (see Methods). Models performed well for the aging human skin dataset [28] (Figure **4**c,d), both when applying model 1 to predict testing data ages (Pearson’s *r*=0.934, *MAE*=2.8 years) and when applying model 2 to predict ages of the original training data (Pearson’s *r*=0.831, *MAE*=4.4 years) using *Ŷ_te_* instead of *Y_te_* to fit model 2. Therefore, we inferred that the training and the testing datasets are interchangeable and can be used reciprocally to predict each other, which is expected for a dataset split randomly into two parts. Similarly, we obtained good reciprocal predictions for the mouse blood samples subset [31] (Figure **S3**e)

In accordance with the already demonstrated evidence that the datasets of aging mouse liver [31, 32] exhibit no significant covariate shift (Figure **S1**d), we observed that they both pass the ITTP relatively well (Figure **S3**b,c). A slightly lower accuracy of predicting Meer ages with clocks trained on Thompson data (Pearson’s *r*=0.737, *MAE*=6.14 months) might be attributed to the presence of a number of outliers in the Meer dataset (Figure **S1**d). At the second step, however, we obtained great performance metrics (Pearson’s *r*=0.975, *MAE*=1.49 months), which is another evidence of absence of a significant covariate shift between these datasets.

Case 2 applications of the ITTP demonstrated more diverse results. Both in vitro fibroblast reprogramming datasets (Figure **4**e and Figure **S3**a) yielded poor predictions for the aging human skin data (Pearson’s *r*=0.235, *MAE*=52.7 years and Pearson’s *r*=0.239, *MAE*=34.3 years for the second-step models trained on the Ohnuki et al. [30] and the Gill et al. [4] datasets, respectively). We therefore concluded that the reprogramming datasets cannot be used to predict normal aging skin data if we consider age predictions for the reprogramming data to be accurate.

The dataset of in vivo reprogramming in mouse liver [3], on the other hand, passed the second step of ITTP with remarkable success, demonstrating the performance metrics of Pearson’s *r*=0.968 and *MAE*=2.21 months (Figure **S3**d). Of note, the predictions at step 1 are shifted upward, while the aging trend is still captured well (Pearson’s *r*=0.826). Finally, applying the ITTP to the datasets of aging mouse blood [31] and mouse embryogenesis [34] resulted in prediction failures similar to that of in vitro reprogramming (Figure **S3**f).

To summarize, the ITTP method highlights the concerns of applying aging clocks to the reprogramming data (or to any other OOD scenario). As an empirical test to discover possible dataset shift, it helps to assess the risk of prediction failure. The results presented above showed that normal aging cannot be predicted using in vitro reprogramming data, which immediately prompts to inquire whether, in return, the ages across reprogramming can be predicted correctly using data on normal aging.

### Uncertainty-aware clocks reveal insignificance of age reversal

In clinical practice, where decision-making often relies on the level of uncertainty, an ML model is required to estimate not only the desired outcome, but also the uncertainty of its predictions [15]. A robust model should be able to warn of extreme uncertainty when making predictions on shifted datasets. The majority of aging clock papers, inspired by Horvath [26] and Hannum et al. [27], have adopted the ElasticNet model that lacks inherent uncertainty estimation. To address this limitation, we turned to the Gaussian Process Regressor (GPR) model [43, 44], a variant of which was recently employed in aging clocks [45] (see Methods).

To demonstrate the concept of GPR, we trained and tested it on a single CpG site (Figure **5**a,b). A Gaussian process (GP) can be viewed as a probability distribution encompassing all possible functions that can be fitted over the training observations [43]. Therefore, for every input methylation value, a fitted GPR model provides an estimation of the most probable age, along with the probability distribution around this estimation with a finite variance that represents a credible interval for the prediction. The credible interval, grounded in Bayesian statistics, is calculated for each individual prediction relying on the training data and prior model assumptions. It should not be confused with a confidence interval which describes only the distribution of multiple predictions.

**Figure 5:**
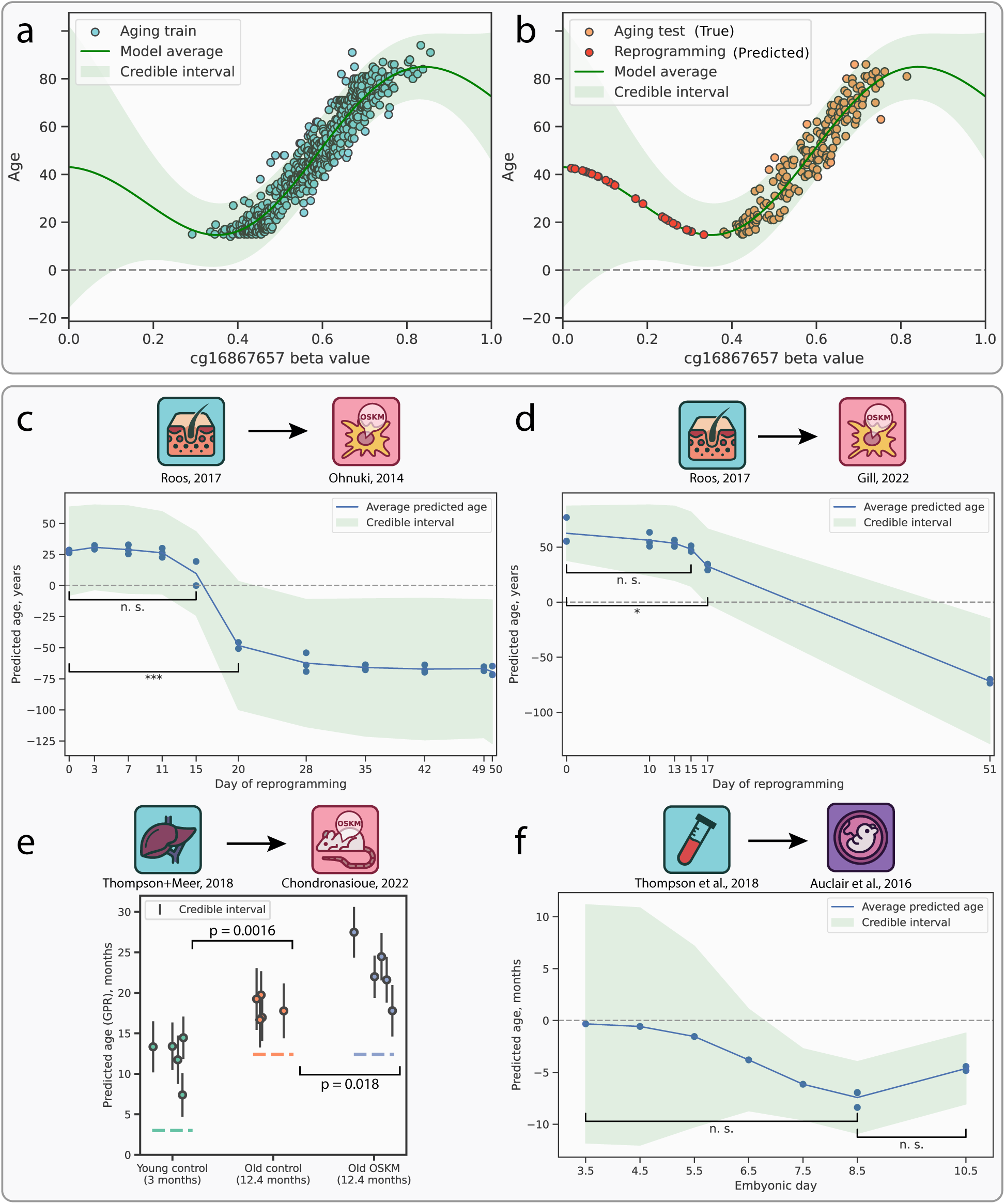
Estimation of epistemic uncertainty in the rejuvenation datasets using Gaussian Process Regression (GPR) models. **a**, A simplified case of a GPR model trained to predict chronological age based on a single methylation site (see Methods). Prediction uncertainty is presented as credible intervals of two standard deviations here and in panels **b-f**. **b**, Prediction uncertainty increases as DNAm values move away from the training distribution. **c-d**, Predicted rejuvenation trajectory and estimated prediction uncertainty for in vitro human fibroblast reprogramming datasets by Ohnuki et al. [30] **(c)** and Gill et al. [4] **(d)**. No significant rejuvenation events are detected before days 20 and 17 of reprogramming, respectively. **e-f**, Predicted epigenetic age trajectory and prediction uncertainty for in vivo mouse reprogramming [3] **(e)** and mouse embryonic development [34] **(f)**.

When a GPR fits the training data well, it computes a finite error stemming from the variation of age points associated with any given DNAm value (earlier referred to as aleatoric uncertainty). As methylation values move further away from the training distribution, the model assigns greater epistemic uncertainty. Consequently, the credible interval widens, reflecting that the model is less familiar with this particular range of DNAm values (Figure **5**b). The GPR thus provides an estimation of total prediction uncertainty, encompassing both its aleatoric and epistemic components, and presenting it as a credible interval of several standard deviations for every individual prediction. Consequently, these credible intervals are employed to compute significance of difference between two prediction groups, such as control and treatment samples or samples from two different days of reprogramming, using a meta-regression approach (see Methods). It is important to note that the GPR predictions are influenced by the choice of prior distributions over functions, meaning that the results may vary depending on these underlying assumptions.

In accordance with the previous sections, we employed GPR models trained on aging datasets to predict rejuvenation trajectories in the respective reprogramming scenarios and to determine the corresponding credible intervals (see Methods). When applied to the in vitro reprogramming dataset [30], a skin-trained GPR (Figure **S4**a) predicted a noticeable epigenetic age decline from reprogramming day 11 through day 28 (Figure **5**c), which aligned well with our previous observations obtained with ElasticNet (Figure **3**a) and with the observations made by the authors of said dataset. However, the accompanying credible interval of two standard deviations revealed how extremely uncertain the model is about these predictions, especially in the case of the late reprogramming phase. This uncertainty prompts questions regarding the significance of any rejuvenation effect until the 20th day of reprogramming, a point at which a complete erasure of somatic identity was reported in the original study [30].

Applying GPR to the in vitro fibroblast reprogramming dataset [4] yielded similar results, with two notable differences (Figure **5**d). First, the credible interval at reprogramming day 0 is slightly narrower, suggesting that these samples might be more similar to the training set. Second, the model indicated a significant rejuvenation effect between days 0 and 17 (*P* value=0.014), indicating that rejuvenation might indeed occur at the later stages of the maturation phase. However, the substantial credible interval at day 17 (spanning from 0 to approximately 70 years) restrains the extent to which confident statements on rejuvenation can be made.

We next revisited the in vivo mouse reprogramming dataset [3] using the GPR model trained on aging mouse liver (Figure **S4**b) and observed a significant negative rejuvenation effect between the control and OSKM-treated old mice (Figure **5**e). This outcome highlights the importance of incorporating individual prediction uncertainties to resolve differences between groups that are otherwise undistinguishable by simpler methods such as Lasso regression. However, similar to Lasso-based predictions, we observed a systematic bias in GPR estimations. This bias did not appear to strongly affect credible intervals, which might suggest that GPR underestimates credible intervals in cases of dataset shift, putatively resulting, in this instance, from the comparison of two different mouse strains.

The GPR model, trained on the mouse blood (Figure **S4**c) and applied to predict the dynamics of epigenetic age during mouse embryogenesis [34], displayed a local minimum at embryonic day 8.5 (Figure **5**f), close to the putative ground zero event at E6.5/E7.5 suggested by Kerepesi et al. [7]. However, our GPR predictions are accompanied by large credible intervals, with insignificant age decline between days 3.5 and 8.5 (*P* value=0.54) and subsequent insignificant age increase between days 8.5 and 10.5 (*P* value=0.37). Consequently, this prevents the definitive designation of day 8.5 as the “ground zero” of epigenetic age. Notably, the credible intervals get narrower throughout the course of embryonic development, supporting our earlier observation of a larger shift between the early days of embryogenesis and the aging process (Figure **S1**j).

Our findings indicate that a GPR model, capable of assessing its prediction uncertainty, can effectively detect covariate shifts in a test dataset by assigning elevated uncertainty to samples not represented in the training data.

## Discussion

Epigenetic clocks have been widely used to demonstrate age acceleration and deceleration in a variety of research contexts [12]. However, there is still a reluctance to include clock measurements as endpoints in the clinical longevity intervention trials [46]. In addition to the fact that the existing clocks fail to capture some aging-associated conditions [20], we emphasize that they cannot be easily validated and relied upon, given their inability to estimate prediction uncertainty.

In this study, we present a computational framework for validating the rejuvenation effects predicted by the epigenetic aging clocks. Due to the unavailability of ground truth values for the biological age, the concept of epigenetic age reversal remains speculative because it cannot be validated directly. Therefore, we have to rely on indirect evidence. To address this challenge, our framework encompasses four specific approaches: covariate shift estimation, comparison of different clock models, the Inverse Train-Test Procedure (ITTP), and the prediction uncertainty estimation using a Gaussian Process Regression (GPR) model.

We demonstrate how the presence of covariate shift can distort model performance and show that covariate shift is highly prominent in the DNA methylation (DNAm) data across cellular reprogramming and embryonic development. Moreover, by applying eight different published aging clocks [26, 27, 37–41] to the in vitro reprogramming data, we illustrate that the magnitude of rejuvenation effect achieved by the end of the maturation phase of reprogramming highly depends on the clock model type and the chosen training data. Across the discussed models, the age reversal effect can differ by up to two orders of magnitude, including null and even negative rejuvenation. Our ITTP approach further reveals that in vitro reprogramming datasets cannot robustly predict normal aging, thus additionally challenging the notion that normal aging can accurately predict reprogramming.

A GPR-based aging clock can estimate the uncertainty of its own age predictions in the form of standard deviations, even for samples that do not belong to its training distribution. While reproducing the decrease in average epigenetic age, this clock demonstrates no statistically significant difference between the days 0 and 15 of in vitro reprogramming in one dataset [30] (Figure **5**c), and marginal significance between days 0 and 17 in another dataset [4] (Figure **5**d). However, the accompanying credible interval at day 17 spans approximately 70 years, thereby preventing definitive conclusions regarding the observed rejuvenation effects.

The in vivo mouse liver reprogramming data [3] exhibits a covariate distribution closer to that of normal mouse aging. Moreover, it passes the ITTP test, suggesting that in vivo reprogramming might preserve organismal states better than the in vitro procedure. However, we show that both the Lasso and the GPR clock models surprisingly predict old reprogrammed samples to be either of the same age or even significantly older than the old controls (Figure **5**e). This finding prompts further forays into the nature of processes accompanying in vivo reprogramming, especially in light of a recent study describing impaired liver function and premature death in mice with continued OSKM induction [47].

While testing the “ground zero” hypothesis of epigenetic aging [33], we discover significant covariate shift and ITTP failure for the dataset spanning from early to middle embryonic development [34]. While average GPR-predicted ages recapitulate previously described dynamics of epigenetic age decrease in embryogenesis [7] and yield age zero at embryonic day 8.5, these dynamics are found to be statistically insignificant from the epistemic uncertainty standpoint (Figure **5**f).

We hypothesize that the GPR model assigns such wide credible intervals to both in vitro reprogramming and embryogenesis because the totipotent and pluripotent states are too unfamiliar to a model trained purely on differentiated somatic cells. Thus, we have shown that an aging clock model that performs well within aging datasets will likely fail to reliably predict rejuvenation events not represented in the training data. To decrease this uncertainty, the inclusion of progenitor cells in the training dataset could prove beneficial.

Our work should not be viewed as an attempt to prove or disprove the existence of rejuvenation effects, including the “ground zero” hypothesis of epigenetic aging. We also do not aim to comprehensively cover all available datasets of putative rejuvenation, be it reprogramming, embryogenesis, or other interventions. Instead, by concentrating on the most striking examples, we illustrate that the existing aging clocks that rely on overly optimistic assumptions [11] cannot serve as reliable biomarkers of rejuvenation.

From the machine learning viewpoint, the effect of out-of-distribution samples on the prediction outcome is well-known [48, 49], as a trained model is highly unlikely to perform robustly on data out of the original distribution [50, 51]. Likewise, the good performance on the training and the validation datasets cannot guarantee that the model trained to predict normal aging will reliably predict the rejuvenation effects. Nor could it be used as the reprogramming validation method, especially given the lack of ground truth and the large resulting uncertainty. Yet, these approaches appear to have become common today, which is one practice that we hope will be reconsidered.

Despite the clock drawbacks, a reliable surrogate health measure [52] is still required for the evaluation of longevity drugs and other interventions in clinical trials [53]. Several criteria have been proposed for the potential biomarkers of aging [54]. We suggest that, in order to gain a wider acceptance within the longevity community, the single-point age clock predictions must be accompanied by the uncertainty estimation [15], while acknowledging for the potential limitations of this estimation. Furthermore, we recommend assessing potential covariate shifts between datasets with methods discussed in this work before applying any aging clock models.

We advocate for the development of a clinically relevant and reliable aging clock with a clearly defined training target, such as mortality, combined with the ability to estimate prediction uncertainties. This approach is essential for alerting researchers to possible misinterpretations of the trial results. While such a protocol may add complexity to aging clock development, it is crucial for advancing the field and for positioning aging clocks as accurate health estimators.

## Materials and Methods

### Height and weight data processing

For our toy example of the dataset shift, we sourced data from the WHO [21] for the control cohort and from Hoover et al. [23] for the achondroplasia cohort. The datasets included height and weight means and standard deviations across various ages: 60–228 months for the control and 0–240+ months for the achondroplasia cohort. We interpolated the control dataset using the methodology by Andres et al. [55] to align age ranges (0–240+ months). Assuming a normal joint distribution for height and weight [22], we sampled points with the corresponding means and covariance matrices. Age was uniformly sampled from 0 to 276 months. 1000 samples were generated for each cohort.

Using 1000 samples from the control cohort, we constructed a bivariate linear regression model with the Python *scikit-learn* [56] package. Post-training, this model was used for age prediction in the achondroplasia cohort.

### DNA methylation data processing

For in vitro reprogramming, we employed two datasets of DNA methylation (DNAm) profiling throughout the human fibroblast reprogramming timeline by Ohnuki et al. [30] and Gill et al. [4], which are the only existing open access DNAm datasets of this kind, and compared them with the aging human skin datasets by Roos et al. [28] and Vandiver et al. [29]. To compare mouse DNAm profiles across aging, in vivo reprogramming, and embryogenesis using as wide genomic CpG coverage as currently possible, we focused on the methylation data obtained by reduced representation bisulfite sequencing (RRBS) [57]. Thus, we employed multi-tissue datasets of mouse aging from Thompson et al. [31] and Meer et al. [32], the in vivo reprogramming data from Chondronasiou et al. [3], and the mouse embryogenesis DNAm dataset by Auclair et al. [34] which stands out as one of the few bulk-tissue DNAm datasets featuring both early and middle embryonic stages.

For the studies of Roos et al. [28] (GSE90124), Vandiver et al. [29] (GSE51954), Ohnuki et al. [30] (GSE54848), Gill et al. [4] (GSE165179), Auclair et al. [34] (GSE60334), and Chondronasiou et al. [3] (GSE156557), files containing processed methylation beta values were downloaded from the GEO database under the corresponding accession numbers. For detailed description of sample groups isolated for our analysis, see Supplementary Table 1. Methylation matrices were assembled for each dataset by merging selected samples and retaining only those CpG sites that appeared in every sample of the respective dataset.

A processed multi-tissue RRBS dataset from the Thompson et al. study was downloaded from the GEO database, where it is deposited under accession no. GSE120132 (ref. [31]). As all other mouse DNAm studies referenced in this study used wild-type C57Bl/6J or related mouse strains, we selected 196 samples representing this strain from the 549 samples comprising the original dataset. In the original dataset, CpG methylation values from both DNA strands were combined and assigned to the forward strand cytosines, while retaining strand annotation. To utilize as much information as possible, we treated methylation values from forward and backward strands separately. A processed multi-tissue RRBS dataset from the Meer et al. study comprising 81 samples was also downloaded from the GEO database under accession no. GSE121141 (ref. [32]).

For the studies of Thompson et al. and Meer et al., we followed the instructions on data processing provided by Trapp et al. [8]: in each dataset, we retained only autosomal DNAm sites that featured at least 5-fold coverage in no less than 90% of samples, which yielded 1,242,194 and 2,131,404 sites, respectively. For the analysis of embryonic development, we further isolated 50 blood samples from the Thompson et al. dataset. For the analysis of in vivo liver reprogramming, we isolated 30 and 20 liver samples from the Thompson et al. and the Meer et al. datasets, respectively, and merged them into one methylation matrix.

### Principles of CpG selection

CpG site selection for aging clock training is a nuanced challenge, with various authors suggesting distinct, minimally overlapping subsets [35]. To tackle this problem, we used CpG sites (CpGs) from established clocks most relevant to each corresponding dataset pair, considering tissue composition of the respective datasets. For instance, in analyzing covariate shift between the aging human skin and in vitro human fibroblast reprogramming datasets, Horvath’s skin clocks [37] and the corresponding CpGs were employed. While any accurate age-predicting CpG subset could suffice, we predominantly used known subsets for methodological simplicity where possible. The detailed description of dataset pairs, clock models, and the amount of clock sites observed in the datasets is specified in Supplementary Table 2.

### Principal component analysis for covariate shift visualization

Principal component analysis (PCA) presented in Figure **2**c,f,i and Figure **S1**a,d,g,j was conducted on merged dataset pairs to select CpGs for the covariate shift analysis (refer to previous sections). This analysis employed the Python *scikit-learn* library [56] with the default parameters. The two-dimensional kernel density estimations of PCA results were performed using the *kdeplot* function from the Python *seaborn* package.

### Kolmogorov-Smirnov test for a shift of individual covariates

We employed a two-tailed Kolmogorov-Smirnov (KS) test on the selected CpGs (refer to site selection principles above) to detect covariate shifts between the dataset pairs. This test assessed the null hypothesis that the beta value distributions for a specific CpG site are identical across a given dataset pair. We applied the Benjamini-Hochberg correction to the computed *P* values, considering the adjusted *P* value below 0.01 as indicative of a significant distributional shift. The KS statistics and the *P* values were calculated using *scipy* [58], and the multiple testing correction was performed with *statsmodels* [59] in Python.

### Testing in vitro reprogramming datasets with different published epigenetic aging clocks

We evaluated the consistency of predictions across eight aging clock models using the R *methylclock*[60] package, which predicts epigenetic age from the input matrices of CpG methylation beta values. The clocks included: Horvath’s multi-tissue clock [26], Hannum’s blood clock [27], Horvath’s skin+blood clock [37] (notated as Horvath skin in this paper), PedBE clock [39], Wu’s clock [40], Zhang’s Best Linear Unbiased Prediction (BLUP) and ElasticNet-based (notated as Zhang in this paper) clocks [41], and Levine’s blood clock (also known as PhenoAge) [38]. All *methylclock*-generated predictions are available in our GitHub repository provided further below.

### Testing in vitro reprogramming datasets with different models of aging clocks trained on the same dataset

To evaluate prediction consistency across different aging clock model families, we employed five different machine learning models from the *scikit-learn* package: *k*-neighbors regressor (the *KNeighborsRegressor* function), random forest regressor (the *RandomForestRegressor* function), support vector regressor (the *SVR* function), Bayesian ridge regressor (the *BayesianRidge* function), and histogram-based gradient boosting regressor (the *HistGradientBoostingRegressor* function). These models were trained on the aging human skin dataset [28] using 5-fold cross-validation and hyperparameter optimization via grid search, assessing model performance with the mean squared error metric. Performance metrics of the optimally-tuned models for both the training and testing subsets are summarized in Table 1.

**Table 1:** Model Performance Comparison.

| Model | $r$ Train | $r$ Test | $R^2$ Train | $R^2$ Test | $MAE$ Train | $MAE$ Test |
| --- | --- | --- | --- | --- | --- | --- |
| $k$-neighbors regressor | 0.844 | 0.654 | 0.699 | 0.402 | 3.915 | 6.037 |
| Random forest regressor | 0.980 | 0.847 | 0.934 | 0.650 | 1.793 | 4.585 |
| Support vector regressor | 0.963 | 0.933 | 0.921 | 0.854 | 1.603 | 2.923 |
| Bayesian ridge regressor | 0.986 | 0.932 | 0.971 | 0.863 | 1.235 | 2.750 |
| Histogram-based gradient boosting regressor | 0.999 | 0.883 | 0.998 | 0.752 | 0.237 | 3.729 |

### Lasso clocks for the in vivo reprogramming testing

We observed that the in vivo reprogramming dataset [3] exhibited limited overlap with three established mouse aging clocks (7/582 sites in the Thompson clocks [31], 3/90 sites in the Meer clocks [32], and 16/436 sites in the Petkovich clocks [61]), potentially impairing clock predictions. Consequently, we developed a new clock using a Lasso penalized regression model, trained on the liver samples combined from the Thompson et al. [31] and Meer et al. [32] aging mouse liver datasets. Utilizing the *LassoCV* class from *scikit-learn* [56], we identified the optimal regularization hyperparameter *α* through 5-fold cross-validation. The final model, selecting 22 out of 16,849 CpGs (Supplementary Table 3), exhibited strong performance while testing (*MAE*=2.2 months, *R*^2^=0.866), as detailed in Figure **3**e. This model was then applied to predict epigenetic age in the in vivo reprogramming dataset [3].

### Inverse Train-Test procedure with Lasso regression

In general, the Inverse Train-Test procedure (ITTP) can be applied to any pair of datasets to test their interchangeability. However, in practice, the outcome of the procedure will depend on the generalizing abilities of the chosen model. Thus, it is crucial to use models from the same family (for instance, linear models) at steps 1 and 2 of the ITTP. We decided to choose a linear regression model with Lasso penalization as a base model (*i.e.*, model 1 and model 2 in Figure **4**a) for the ITTP because it has generalization properties equivalent to ElasticNet (most often used aging clock model), but is easier to train (optimizing one hyperparameter instead of two). Below, we provide a detailed algorithm for the ITTP procedure for both use cases discussed in this study, presuming that the Lasso model is used as the base model.

### ITTP case 1

The ITTP case 1 (also referred to as the verifiable case) considers a pair of datasets having ground truth values: the training dataset {*X_tr_, Y_tr_*} and the testing dataset {*X_te_, Y_te_*}, where *X_tr_* and *X_te_* denote training and testing features (CpG methylation values), respectively, and *Y_tr_* and *Y_te_* denote the corresponding training and testing targets (chronological ages). Let two different initializations of the Lasso regression model be 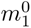 and 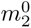for the corresponding steps 1 and 2 of ITTP, where the superscript 0 denotes the model state before training. Accordingly, let 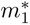 and 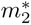 denote the models after training. Then, the ITTP case 1 can be performed by the following algorithm:

#### Step 1

1. Train model 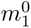 on {*X_tr_, Y_tr_*}. Select the optimal regularization parameter *α* for Lasso regression, employing cross-validation to evaluate model performance across a range of alpha values [*α_min_, α_max_*]. For that, split the dataset {*X_tr_, Y_tr_*} into multiple training and validation sets (we used 5-fold splitting), train the model on each, and assess performance using the mean squared error metric. The *α* value yielding the best average performance across all folds is chosen as the optimal one.
2. Apply the trained model 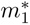 to test dataset *X_te_* predicting *Ŷ_te_*.
3. Compute performance metrics for the model 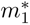 predictions for the pair *Ŷ_te_* and *Y_te_*. We propose to compute widely used regression metrics such as *R*^2^, mean absolute error (*MAE*), and Pearson’s correlation coefficient (*r*).

#### Step 2

1. Train model 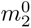 on {*X_te_, Ŷ_te_*}. Select the optimal regularization parameter *α*, as in Step 1.
2. Apply the trained model 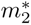 to train dataset *X_tr_* predicting *Ŷ_tr_*.
3. Compute performance metrics (*R*^2^, *MAE*, *r*) for the model 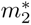 predictions by comparing *Ŷ_tr_* and *Y_tr_*.

Compare the performance metrics obtained from the steps 1 and 2 of the ITTP. If the metrics are satisfactory and comparable, then the datasets are assumed to be interchangeable.

Within the study, a threshold of *R*^2^>0.25 was employed to ascertain whether a model exhibits satisfactory performance. If satisfactory metrics are obtained only during step 2, then we can assume that the given datasets are not fully interchangeable, but the testing dataset still contains the information required for the training dataset prediction.

### ITTP case 2

The ITTP case 2 (also referred to as the unverifiable case) considers a pair of datasets where the ground truth values are available for the training dataset only: the training dataset {*X_tr_, Y_tr_*} and the testing dataset {*X_rep_*} (we denoted the testing dataset as *X_rep_* to emphasize that the reprogramming datasets do not contain the ground truth values of biological age). The algorithm for this case is similar to the ITTP case 1 with some distinctions in step 1:

#### Step 1

1. Train model 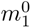 on {*X_tr_, Y_tr_*}. Select the optimal regularization parameter *α*, as was described in the paragraph regarding step 1 of the ITTP case 1.
2. Apply the trained model 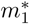 to the testing dataset *X_rep_* to yield age predictions *Ŷ_rep_*.

#### Step 2

1. Train model 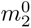 on {*X_rep_, Ŷ_rep_*}. Select the optimal regularization parameter *α*, as was described in the paragraph regarding step 1 of the ITTP case 1.
2. Apply the trained model 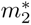 to the training dataset *X_tr_* to yield age predictions *Ŷ_tr_*.
3. Compute performance metrics (*R*^2^, *MAE*, *r*) for the model 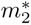 predictions by comparing *Ŷ_tr_* and *Y_tr_*.

In the second case of ITTP, we rely only on the performance metrics computed during step 2. If the metrics are satisfactory, the datasets can be assumed interchangeable. Otherwise, the datasets cannot be used to predict each other according to the chosen linear model assumption.

To train Lasso models during the ITTP steps described above, we used the *LassoCV* class from the Python *scikit-learn* library [56], which conducts a search for the optimal Lasso regularization hyperparameter *α* with inbuilt cross-validation over the training subset.

### Prediction uncertainty inference using Gaussian Process regression model

A Gaussian Process regression (GPR) model was developed to predict the age of samples given their methylome. GPR is a flexible non-parametric Bayesian approach to the regression tasks [43]. In our model, the inputs are the same CpG sites used for covariate shift analysis from the published and the newly constructed (see the section about Lasso training) aging clocks (Supplementary Table 2), and the outputs are the ages of the sample donors. The model was trained using the Python *scikit-learn* package with a composite kernel comprising the Radial Basis Function (RBF) and the white noise kernels elaborated further below.

Given that the methodology we use to build the GPR-based aging clocks has been described in detail elsewhere [45], this section is focused primarily on deriving the prediction uncertainty, which is essential for our study. A Gaussian Process (GP) is a probability distribution over possible functions that fit a set of points. Formally, it is a collection of random variables, any finite number of which have a joint Gaussian distribution [43]. Given a train sample set 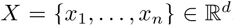, a mean function 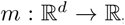, and a covariance function 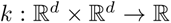, a GP can be written as 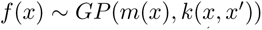, if the outputs *f* = (*f* (*x*_1_)*, … , f* (*x_n_*))*^T^* have a Gaussian distribution described by 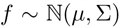, where *µ* = *m*(*x*_1_*, … , x_n_*) and Σ*_i,j_* = *k*(*x_i_, x_j_*). The mean function is usually assumed to be the zero function, and the covariance function is a kernel function chosen based on assumptions about the function to be modeled. We tried different kernel functions and found that the sum of the Radial Basis Function and the white noise kernels performed the best in terms of prediction metrics (*MAE* and *R*^2^). The RBF kernel was also used by the authors of the previously reported GPR-based aging clock [45]. The RBF kernel is defined as:

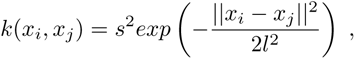

where *s*^2^ is the variance hyperparameter and *l* is the length-scale hyperparameter to control the smoothness of the modeled function, or the speed of its variation. Because GP assumes the output variable includes an additive Gaussian noise part 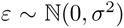, i.e. *y_i_* = *f* (*x_i_*) + *ε*, the vector of outputs is viewed as 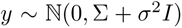. The term *σ*^2^*I* reflects the white noise kernel added to the model to account for the noise in observations, which corresponds to the aleatoric part of the prediction uncertainty.

Given a test point *x^∗^*, its output distribution is defined as *f ^∗^|x^∗^, X, y* which is a conditional Gaussian distribution of the following form:

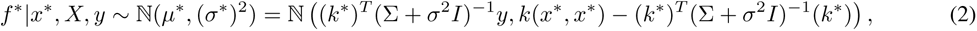

where *k^∗^* = (*k*(*x*_1_*, x^∗^*)*, … , k*(*x_n_, x^∗^*))*^T^* .

Thus, given the training data, the distribution of predictions of a new point is given by a closed analytical form of Gaussian distribution. In our model, the inputs are the CpG methylation vectors, and the outputs are the chronological ages. The mean of the distribution *µ^∗^* can be used as the final prediction of the regression model (corresponds to the prediction of ElasticNet or other models). At the same time, the variance of the distribution (*σ^∗^*)^2^ expresses the level of total prediction uncertainty — one of the most important aspects of our study. One can see that the magnitude of the uncertainty depends on the vector of covariates *k^∗^* of the new sample *x^∗^* with the training set samples *X*. Because the RBF kernel relies on the quadratic distance between the samples, the total prediction uncertainty for an OOD sample will increase as the sample moves away from the training distribution until it reaches the limiting value 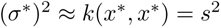, determining the upper bound of the prediction uncertainty the model can estimate for an OOD sample.

### Testing the rejuvenation effects using a meta-regression approach

The Gaussian process model, yielding uncertainty levels for individual sample predictions as Gaussian distribution variances, enables statistical comparison of two predictions via, for example, the *z*-test (if two variances are assumed to be equal). For comparing prediction groups, each with unique variances, we employed advanced meta-analysis techniques. Utilizing the *meta_regression* function from Python *PyMARE* 0.0.3 [62], we accounted for individual age prediction variances in two in vitro reprogramming groups (*e.g.*, days 0 and 15). This function, which incorporates average ages, variances, and a binary group indicator in the design matrix, uses a restricted maximum likelihood approach to optimize meta-regression coefficients, providing coefficient estimates and their *P* values.

### Computational and statistical analyses

Except for the *methylclock* 0.7.8 predictions conducted via R 4.2, all other analyses were performed in Python 3.9.18 using *numpy* 1.22.4, *pandas* 1.5.1, *scipy* 1.7.2, and *scikit-learn* 1.2.1 for data handling and the majority of calculations, and *matplotlib* 3.5.1 in combination with *seaborn* 0.11.2 for data visualization. Multiple testing corrections were performed with *statsmodels* 0.13.2 where indicated. Other packages and functions used for specific computational and statistical analyses are cited in the corresponding sections above.

## Supporting information

Supplemetary Table 1 - Utilized datasets

Supplemetary Table 2 - Train-Test pairs of datasets

Supplemetary Table 3 - Lasso clock coefficients

## Data availability

All datasets of DNA methylation (DNAm) in this study were obtained from the Gene Expression Omnibus (GEO) under the corresponding accession numbers (GSE IDs). Dataset details including GEO IDs are available in Supplementary Table 1. The processed versions of these datasets were made accessible via our GitHub repository at https://github. com/ComputationalAgingLab/reprogramming_ood.

## Code availability

The complete code used for all analyses carried out in this study and the detailed installation instructions are publicly available from https://github.com/ComputationalAgingLab/reprogramming_ood.

## Acknowledgments

We would like to thank Leonid Peshkin for substantive discussions in the early stages of preparing this work. Additionally, we thank Mikhail Gelfand for providing valuable suggestions and insightful comments on our manuscript. This work was supported by Program “Skolkovo Institute of Science and Technology – University of Sharjah Joint Projects: Artificial Intelligence for Life”.

## Author contributions

DK and EK conceived the study. DK wrote the code for an essential part of data analysis and prepared the initial version of the figures. EK prepared the final version of all figures and contributed to the human weight-height data analysis. EE prepared DNA methylation datasets and substantially elaborated on the manuscript. DK, EK, and EE prepared the initial version of the manuscript. EEK and DVD supervised the study. All authors contributed to the final version of the article.

---

**Figure S1:**
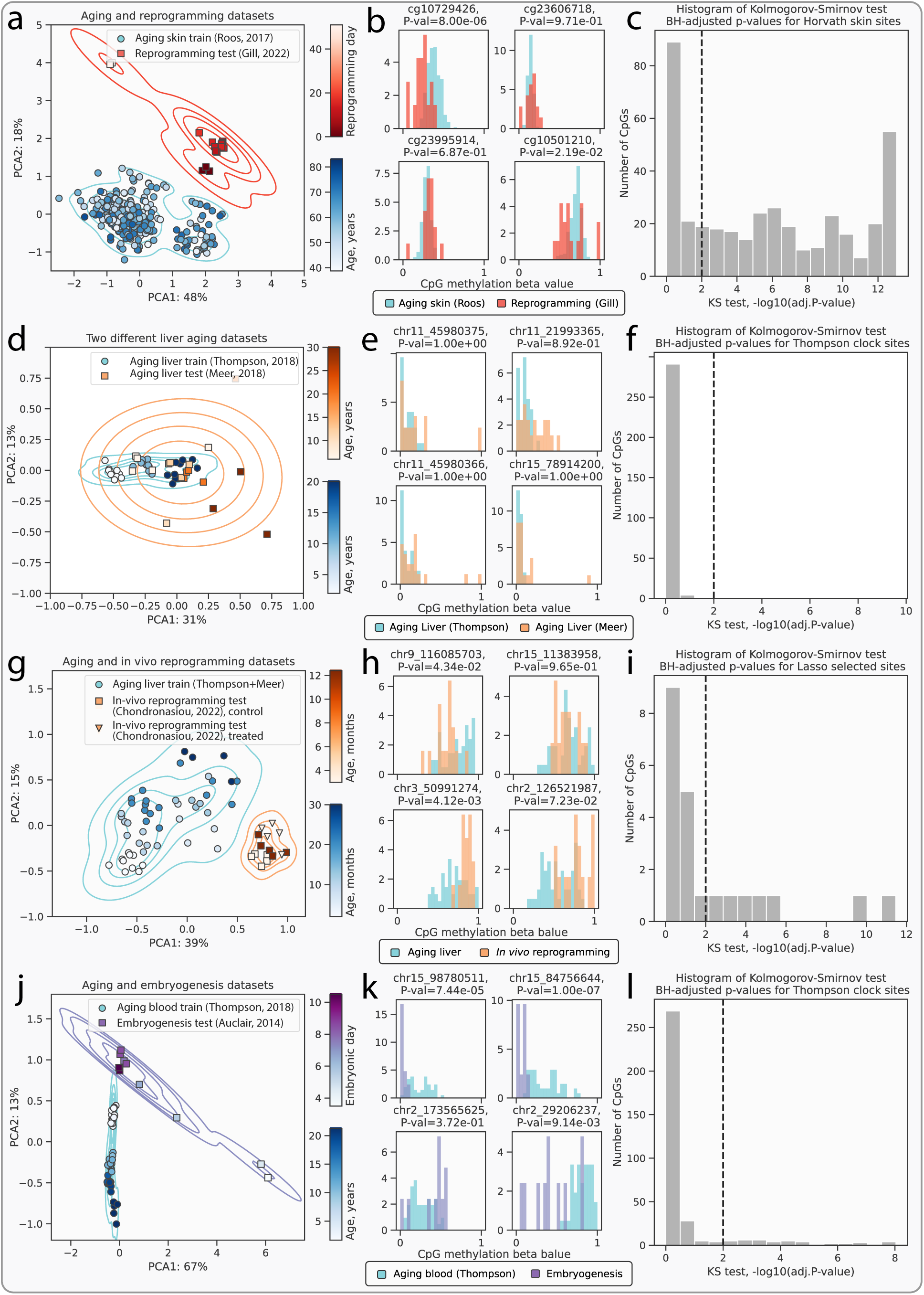
Identification of covariate shift in additional pairs of datasets used in this study. **a-c**, An example of the presence of a substantial covariate shift between the aging skin dataset from Roos et al. [28] and partial *in vitro* fibroblast reprogramming dataset from Gill et al. [4]. **a**, Principal component analysis (PCA) shows a heavy covariate shift between the aging skin and the reprogramming datasets. **b**, Histograms of beta values at top-4 age-correlated CpG sites demonstrate moderate shifts between the aging skin and the reprogramming datasets. **c**, Histogram of the *−log*_10_(*adj.P − values*) demonstrates that 69% of CpG sites were rejected by the two-sample Kolmogorov-Smirnov (KS) test (see Methods) confirming the presence of covariate shift. **d-f**, An example of negligible covariate shift between different aging liver datasets from the Thompson et al. [31] and Meer et al. [32]. **d**, Principal component analysis (PCA) shows minimal discrepancy between the two aging liver datasets with a number of outliers present in the Meer et al. dataset [32]. **e**, Histograms of beta values at four individual CpG sites demonstrate insignificant shifts between the aging liver samples from two different studies. **f**, Histogram of the *−log*_10_(*adj.P − values*) demonstrates that only 1% of CpG sites were rejected by the two-sample KS test (see Methods) confirming the negligible covariate shift. **g-i**, An example of the presence of a moderate covariate shift between the merged aging liver dataset and the transient *in vivo* mouse reprogramming dataset from Chondronasiou et al. [3]. **g**, Principal component analysis (PCA) reveals a moderate covariate shift between the two datasets. **h**, Histograms of beta values at four individual CpG sites demonstrate moderate shifts between the aging liver and the reprogramming control/treatment datasets. **i**, Histogram of the *−log*_10_(*adj.P − values*) demonstrates that 32% CpG sites were rejected by the two-sample KS test (see Methods) confirming the presence of a moderate covariate shift. **j-l**, An example of the presence of a substantial covariate shift between the aging mouse blood dataset from Thompson et al. [31] and the mouse embryogenesis dataset from Auclair et al. [34]. **j**, Principal component analysis (PCA) shows a heavy covariate shift between the aging and the embryonic datasets. Particularly, the data points from the early stages of embryogenesis are significantly divergent from those of the aging blood samples. Conversely, data from the later stages of embryonic development exhibit a closer alignment with the young blood samples. **k**, Histograms of beta values at four individual CpG sites demonstrate varying shifts between the aging blood and the embryonic datasets. **l**, In contrast to the PC analysis (**j**), the histogram of the *−log*_10_(*adj.P − values*) demonstrates that only 12% CpG sites were rejected by the two-sample KS test (see Methods) suggesting a moderate covariate shift. Percents on the axes in **c**, **f**, **i** demonstrate the amount of variance explained by the corresponding principal components. CpG sites for histograms **d**, **g**, **j** were chosen based from the top-four age-correlated sites.

**Figure S2:**
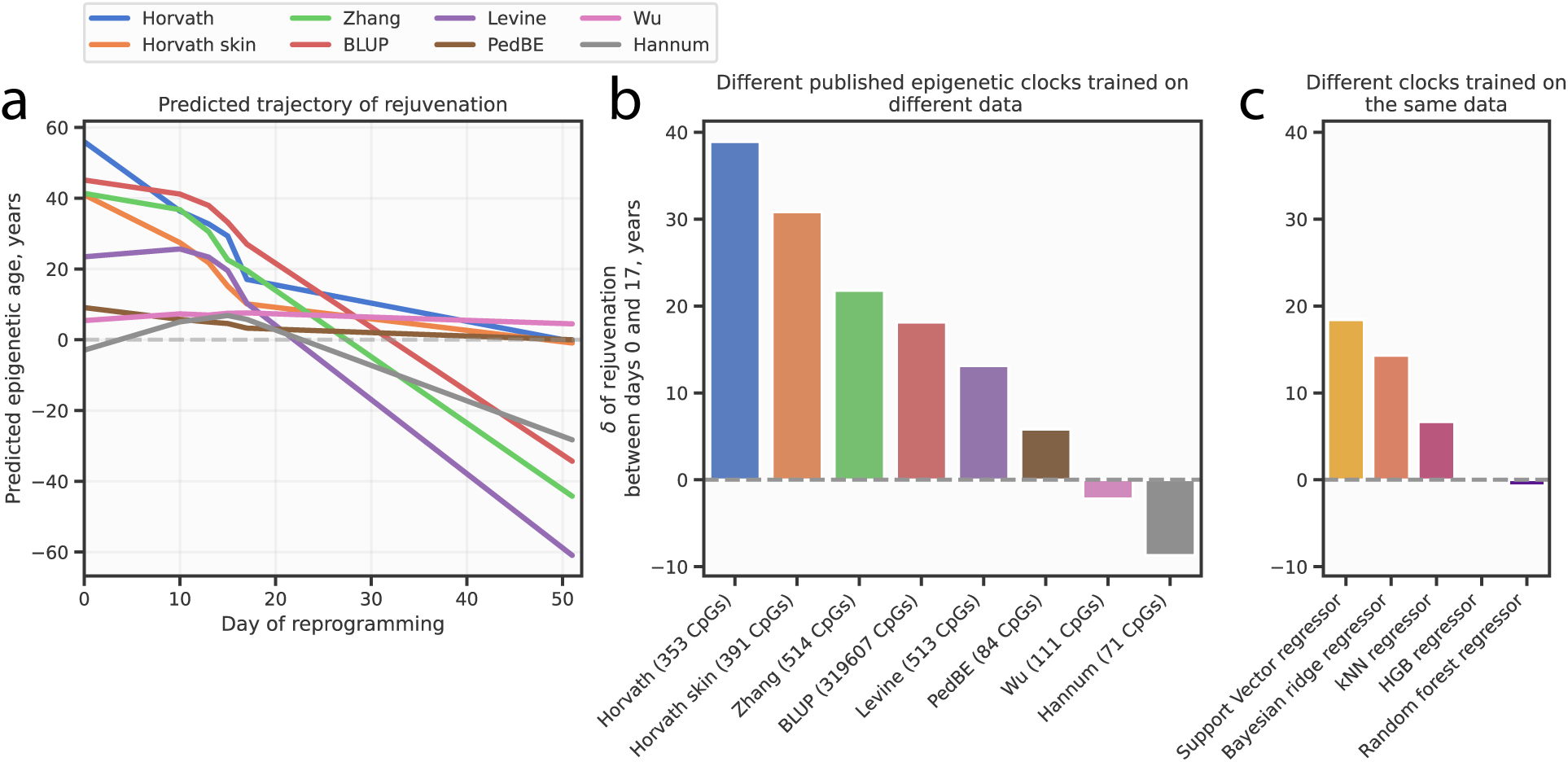
Aging clocks demonstrate inconsistency in the prediction of rejuvenation effect in an additional dataset. **a**, For the Gill et al. [4] dataset, different published aging clocks predict diverse trajectories of rejuvenation during the reprogramming process, which is a manifestation of model uncertainty. Most clocks are ElasticNet models trained on different DNAm datasets. **b**, Aging clocks show differences in accumulated rejuvenation effects between the reprogramming days 0 and 17 calculated with respect to **a**. **c**, Prediction inconsistency holds for the clocks trained on the same aging dataset using different model types, which is another manifestation of model uncertainty.

**Figure S3:**
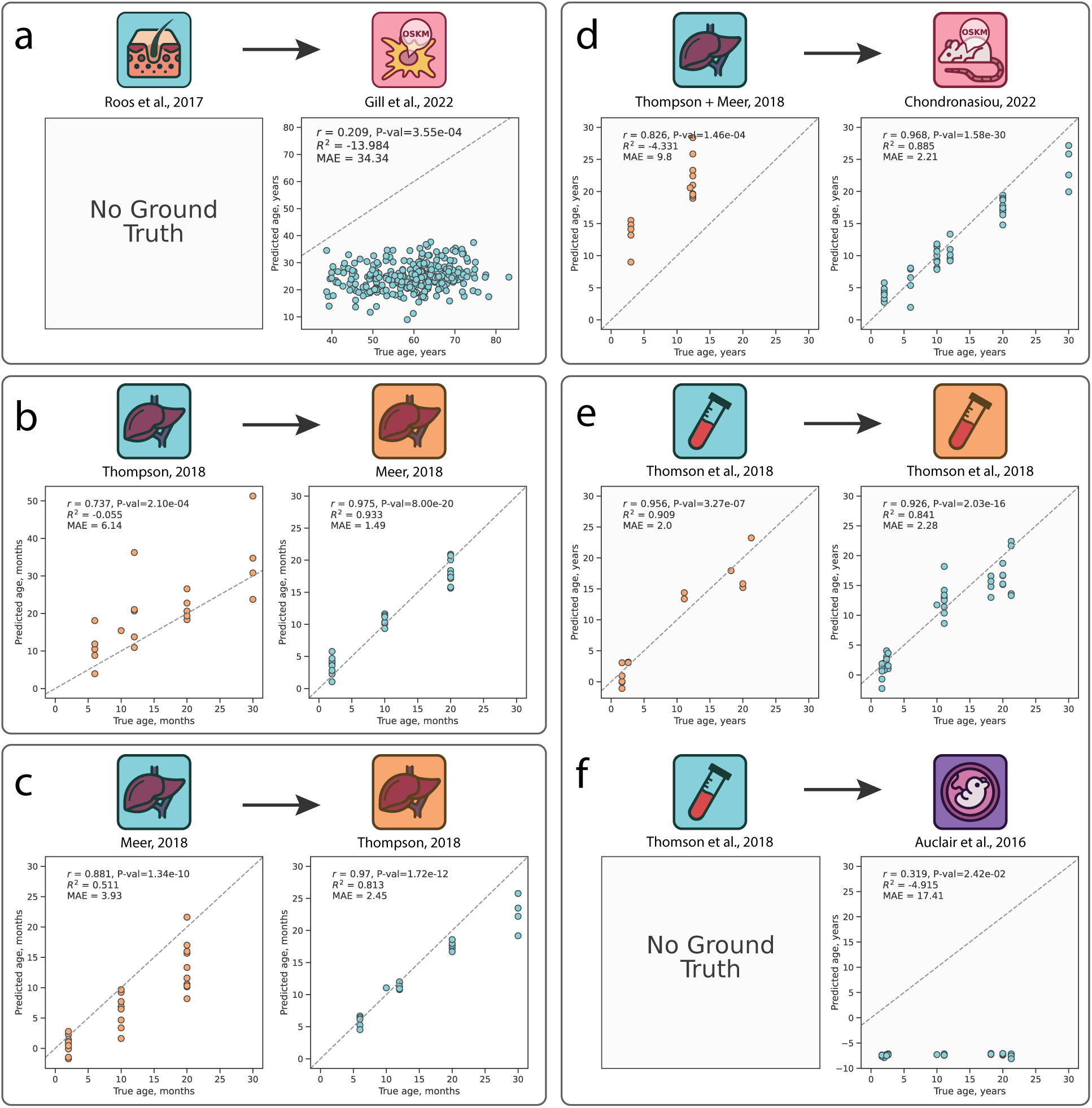
Inverse Train-Test Procedure (ITTP) applied to additional pairs of datasets in the study. **a**, Application of the ITTP to the aging skin dataset [28] and the reprogramming dataset [4]. Poor performance metrics in the second step qualify datasets as non-interchangeable. **b**, Application of the ITTP to two aging mouse liver datasets [31, 32], where the Thompson et al. samples are used for training and the Meer et al. samples are used for testing. The performance metrics are good in both steps, therefore this pair of datasets can be assumed interchangeable. **c**, Inversion of the training and testing datasets employed in **b**. The performance metrics are good on both steps, therefore this pair of datasets can be assumed interchangeable as well. **d**, Application of the ITTP to the combined aging mouse liver dataset [28, 32] used for training and the *in vivo* reprogramming dataset [3] used for testing. The performance metrics are good only for the second step, which nevertheless allows using the reprogramming dataset to predict the aging dataset. **e**, Application of the ITTP to the aging mouse blood dataset [31] split into the training and the testing subsets as 75% and 25% correspondingly (see Methods). The performance metrics are good in both cases, therefore this pair of datasets can be assumed interchangeable. **f**, Application of the ITTP to the aging skin dataset [31] used for training and the embryogenesis dataset [34] used for testing. Poor performance metrics in the second step qualify datasets as non-interchangeable. The blue icons indicate the training aging datasets, the orange icons indicate the testing aging datasets, the red icons indicate the reprogramming datasets, and the purple icon indicates the embryonic dataset. Detailed dataset descriptions can be found in Supplementary Tables 1 and 2.

**Figure S4:**
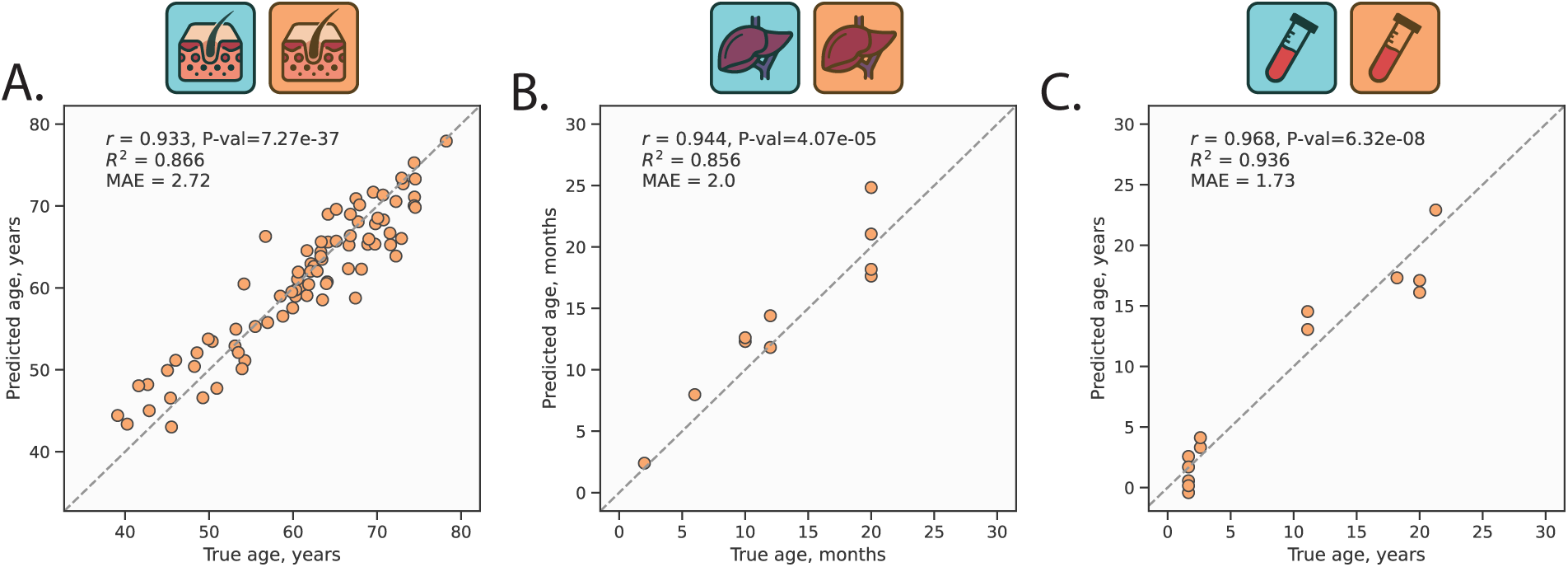
Performance of the GPR models trained on different datasets. **a**, Scatter plot of GPR performance on the testing subset (see Methods). The model was trained on the aging human skin dataset [28]. **b**, Scatter plot of GPR performance on the testing subset (see Methods). The model was trained on a combined aging mouse liver dataset [31, 32]. **c**, Scatter plot of GPR performance on the testing subset (see Methods). The model was trained on the aging mouse blood dataset [31]. Blue icons indicate the training datasets and orange icons indicate the testing datasets. Detailed dataset descriptions can be found in Supplementary Tables 1 and 2.

