## Extended Data Figures for "Epistemic uncertainty challenges aging clock reliability in predicting rejuvenation effects"

Supplementary Materials for

### Epistemic uncertainty challenges aging clocks to predict rejuvenation events

Dmitrii Kriukov, Ekaterina Kuzmina, Evgeniy Efimov, Dmitry V. Dylov, Ekaterina Khrameeva

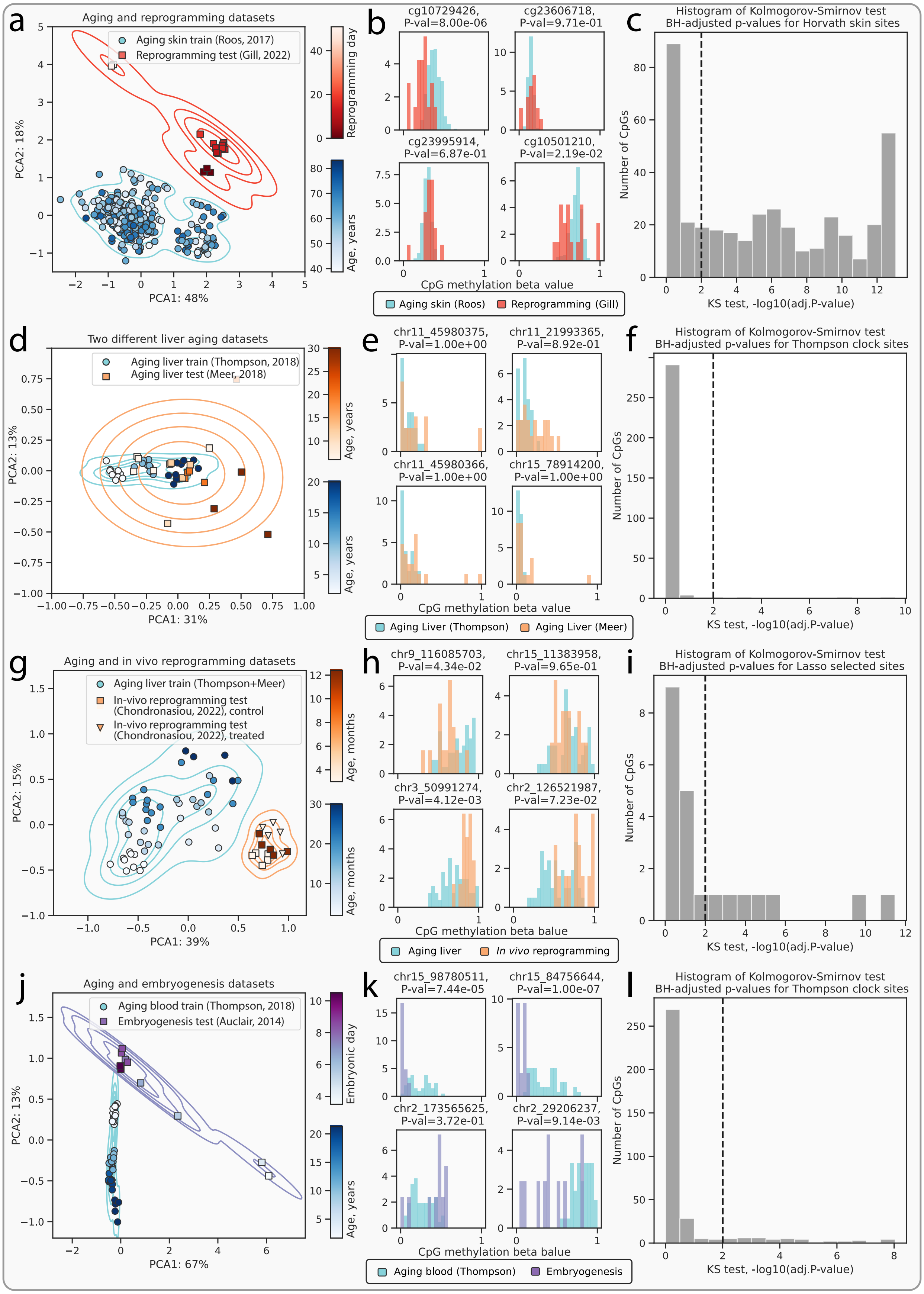

a
Aging and reprogramming datasets
Aging skin train (Roos, 2017)
Reprogramming test (Gill, 2022)
PCA2: 18%
PCA1: 48%
Reprogramming day
Age, years
b
cg10729426, P-val=8.00e-06
cg23606718, P-val=9.71e-01
cg23995914, P-val=6.87e-01
cg10501210, P-val=2.19e-02
CpG methylation beta value
Aging skin (Roos)
Reprogramming (Gill)
c
Histogram of Kolmogorov-Smirnov test BH-adjusted p-values for Horvath skin sites
Number of CpGs
KS test, -log10(adj.P-value)
d
Two different liver aging datasets
Aging liver train (Thompson, 2018)
Aging liver test (Meer, 2018)
PCA2: 13%
PCA1: 31%
Age, years
e
chr11_45980375, P-val=1.00e+00
chr11_21993365, P-val=8.92e-01
chr11_45980366, P-val=1.00e+00
chr15_78914200, P-val=1.00e+00
CpG methylation beta value
Aging Liver (Thompson)
Aging Liver (Meer)
f
Histogram of Kolmogorov-Smirnov test BH-adjusted p-values for Thompson clock sites
Number of CpGs
KS test, -log10(adj.P-value)
g
Aging and in vivo reprogramming datasets
Aging liver train (Thompson+Meer)
In-vivo reprogramming test (Chondronasiou, 2022), control
In-vivo reprogramming test (Chondronasiou, 2022), treated
PCA2: 15%
PCA1: 39%
Age, months
h
chr9_116085703, P-val=4.34e-02
chr15_11383958, P-val=9.65e-01
chr3_50991274, P-val=4.12e-03
chr2_126521987, P-val=7.23e-02
CpG methylation beta balue
Aging liver
In vivo reprogramming
i
Histogram of Kolmogorov-Smirnov test BH-adjusted p-values for Lasso selected sites
Number of CpGs
KS test, -log10(adj.P-value)
j
Aging and embryogenesis datasets
Aging blood train (Thompson, 2018)
Embryogenesis test (Auclair, 2014)
PCA2: 13%
PCA1: 67%
Embryonic day
Age, years
k
chr15_98780511, P-val=7.44e-05
chr15_84756644, P-val=1.00e-07
chr2_173565625, P-val=3.72e-01
chr2_29206237, P-val=9.14e-03
CpG methylation beta balue
Aging blood (Thompson)
Embryogenesis
l
Histogram of Kolmogorov-Smirnov test BH-adjusted p-values for Thompson clock sites
Number of CpGs
KS test, -log10(adj.P-value)

**Extended Data Figure 1**: Identification of covariate shift in additional pairs of datasets used in this study. **a-c**, An example of the presence of a substantial covariate shift between the aging skin dataset from Roos et al. [28] and partial *in vitro* fibroblast reprogramming dataset from Gill et al. [4]. **a**, Principal component analysis (PCA) shows a heavy covariate shift between the aging skin and the reprogramming datasets. **b**, Histograms of beta values at four individual CpG sites demonstrate moderate shifts between the aging skin and the reprogramming datasets. **c**, Histogram of the $-log_{10}(adj.P - values)$ demonstrates that 69% of CpG sites were rejected by the two-sample Kolmogorov-Smirnov (KS) test (see Methods) confirming the presence of covariate shift. **d-f**, An example of negligible covariate shift between different aging liver datasets from the Thompson et al. [31] and Meer et al. [32]. **d**, Principal component analysis (PCA) shows minimal discrepancy between the two aging liver datasets with a number of outliers present in the Meer et al. dataset [32]. **e**, Histograms of beta values at four individual CpG sites demonstrate insignificant shifts between the aging liver samples from two different studies. **f**, Histogram of the $-log_{10}(adj.P - values)$ demonstrates that only 1% of CpG sites were rejected by the two-sample KS test (see Methods) confirming the negligible covariate shift. **g-i**, An example of the presence of a moderate covariate shift between the merged aging liver dataset and the transient *in vivo* mouse reprogramming dataset from Chondronasiou et al. [3]. **g**, Principal component analysis (PCA) reveals a moderate covariate shift between the two datasets. **h**, Histograms of beta values at four individual CpG sites demonstrate moderate shifts between the aging liver and the reprogramming control/treatment datasets. **i**, Histogram of the $-log_{10}(adj.P - values)$ demonstrates that 32% CpG sites were rejected by the two-sample KS test (see Methods) confirming the presence of a moderate covariate shift. **j-l**, An example of the presence of a substantial covariate shift between the aging mouse blood dataset from Thompson et al. [31] and the mouse embryogenesis dataset from Auclair et al. [34]. **j**, Principal component analysis (PCA) shows a heavy covariate shift between the aging and the embryonic datasets. Particularly, the data points from the early stages of embryogenesis are significantly divergent from those of the aging blood samples. Conversely, data from the later stages of embryonic development exhibit a closer alignment with the young blood samples. **k**, Histograms of beta values at four individual CpG sites demonstrate varying shifts between the aging blood and the embryonic datasets. **l**, In contrast to the PC analysis (**j**), the histogram of the $-log_{10}(adj.P - values)$ demonstrates that only 12% CpG sites were rejected by the two-sample KS test (see Methods) suggesting a moderate covariate shift. Percents on the axes in **c**, **f**, **i** demonstrate the amount of variance explained by the corresponding principal components. CpG sites for histograms **d**, **g**, **j** were chosen based from the top-four age-correlated sites.

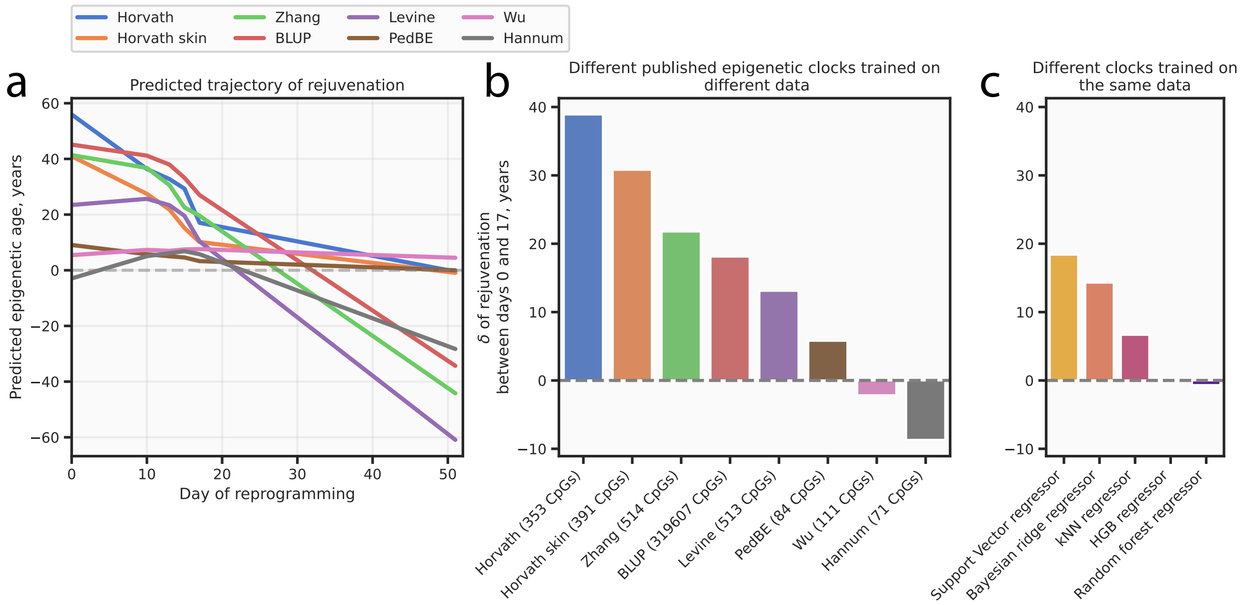


**Extended Data Figure 2**: **Aging clocks demonstrate inconsistency in the prediction of rejuvenation effect in an additional dataset.** **a**, For the Gill et al. [4] dataset, different published aging clocks predict diverse trajectories of rejuvenation during the reprogramming process, which is a manifestation of model uncertainty. Most clocks are ElasticNet models trained on different DNAm datasets. **b**, Aging clocks show differences in accumulated rejuvenation effects between the reprogramming days 0 and 17 calculated with respect to **a**. **c**, Prediction inconsistency holds for the clocks trained on the same aging dataset using different model types, which is another manifestation of model uncertainty.

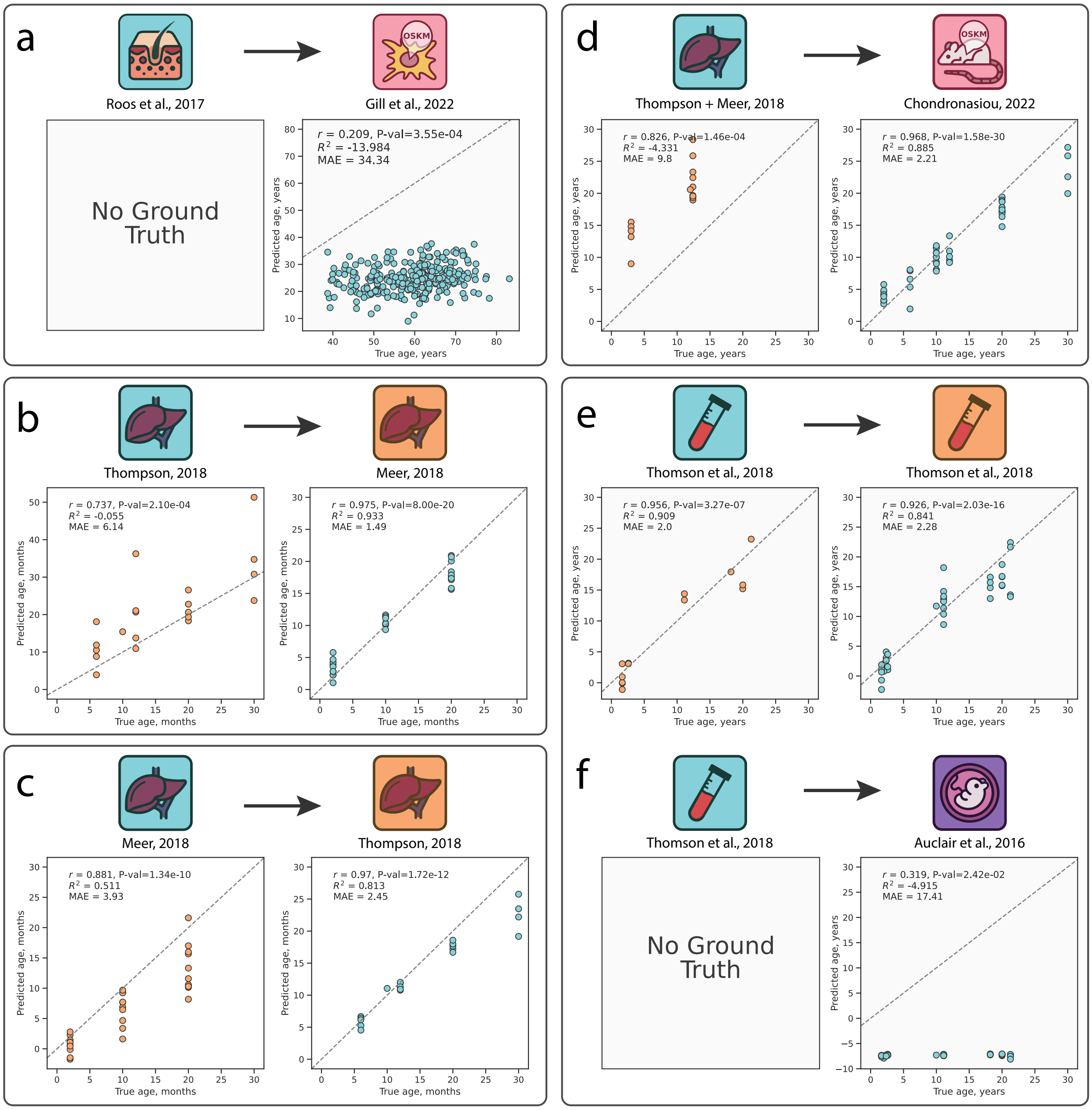


**Extended Data Figure 3**: Inverse Train-Test Procedure (ITTP) applied to additional pairs of datasets in the study. **a**, Application of the ITTP to the aging skin dataset [28] and the reprogramming dataset [4]. Poor performance metrics in the second step qualify datasets as non-interchangeable. **b**, Application of the ITTP to two aging mouse liver datasets [31, 32], where the Thompson et al. samples are used for training and the Meer et al. samples are used for testing. The performance metrics are good in both steps, therefore this pair of datasets can be assumed interchangeable. **c**, Inversion of the training and testing datasets employed in **b**. The performance metrics are good on both steps, therefore this pair of datasets can be assumed interchangeable as well. **d**, Application of the ITTP to the combined aging mouse liver dataset [28, 32] used for training and the *in vivo* reprogramming dataset [3] used for testing. The performance metrics are good only for the second step, which nevertheless allows using the reprogramming dataset to predict the aging dataset. **e**, Application of the ITTP to the aging mouse blood dataset [31] split into the training and the testing subsets as 75% and 25% correspondingly (see Methods). The performance metrics are good in both cases, therefore this pair of datasets can be assumed interchangeable. **f**, Application of the ITTP to the aging skin dataset [31] used for training and the embryogenesis dataset [34] used for testing. Poor performance metrics in the second step qualify datasets as non-interchangeable. The blue icons indicate the training aging datasets, the orange icons indicate the testing aging datasets, the red icons indicate the reprogramming datasets, and the purple icon indicates the embryonic dataset. Detailed dataset descriptions can be found in Supplementary Tables 1 and 2.

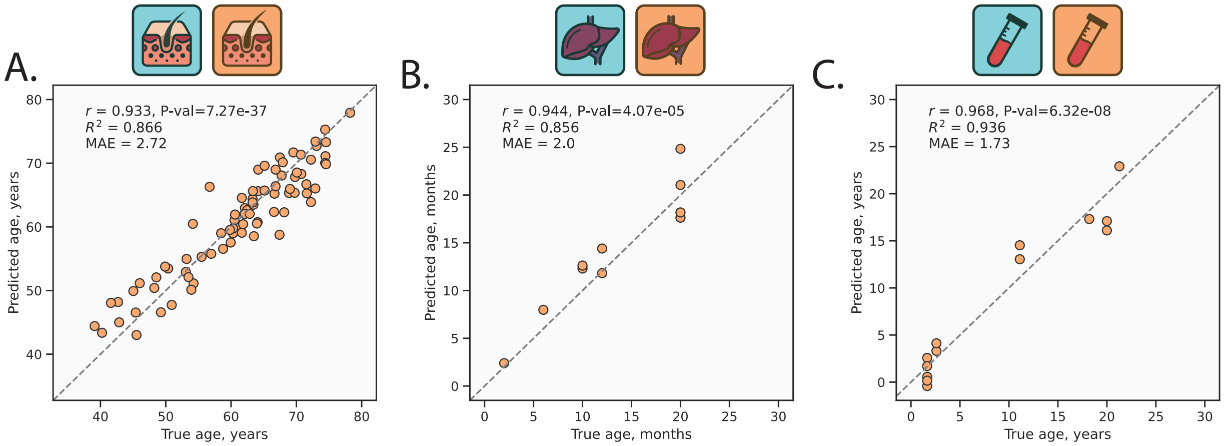


**Extended Data Figure 4**: Performance of the GPR models trained on different datasets. **a**, Scatter plot of GPR performance on the testing subset (see Methods). The model was trained on the aging human skin dataset [28]. **b**, Scatter plot of GPR performance on the testing subset (see Methods). The model was trained on a combined aging mouse liver dataset [31, 32]. **c**, Scatter plot of GPR performance on the testing subset (see Methods). The model was trained on the aging mouse blood dataset [31]. Blue icons indicate the training datasets and orange icons indicate the testing datasets. Detailed dataset descriptions can be found in Supplementary Tables 1 and 2.
